# CORTICOTROPIN-RELEASING FACTOR- DOPAMINE INTERACTIONS IN MALE AND FEMALE MACAQUE: BEYOND THE CLASSIC VTA

**DOI:** 10.1101/2022.04.26.489587

**Authors:** E.A. Kelly, T.M. Love, J.L. Fudge

## Abstract

Dopamine (DA) is involved in stress and stress-related illnesses, including many psychiatric disorders. Corticotropin-releasing factor (CRF) plays a role in stress responses and targets the ventral midbrain DA system which is composed of DA and non-DA cells, and divided into specific subregions. Although CRF inputs to the midline A10 nuclei ('classic VTA') are known, in monkeys, CRF-containing terminals are also highly enriched in the expanded A10 parabrachial pigmented nucleus (PBP) and in the A8 retrorubral field subregions. We characterized CRF-labeled synaptic terminals on DA (tyrosine hydroxylase, TH+) and non-DA (TH-) cell types in the PBP and A8 regions using immuno-reactive electron microscopy (EM) in male and female macaques. CRF labeling was present mostly in axon terminals, which mainly contacted TH-negative dendrites in both subregions. Most CRF-positive terminals had inhibitory (symmetric) profiles. In both PBP and A8, CRF symmetric (inhibitory) synapses onto TH-negative dendrites were significantly greater than asymmetric (excitatory) profiles. This overall pattern was similar in males and females, despite shifts in the size of these effects between regions depending on sex. Because stress and gonadal hormone shifts can influence CRF expression, we also did hormonal assays over a 6-month time period, and found little variability in basal cortisol across similarly housed animals at the same age. Together our findings suggest that at baseline, CRF-positive synaptic terminals in the primate PBP and A8 are poised to regulate DA indirectly through synaptic contacts onto non-DA neurons.

## INTRODUCTION

The neurotransmitter dopamine (DA) is important in many fundamental behaviors including positive and negative reinforcement, decision making, working memory, incentive and stimulus salience and purposeful movement. The majority of DA neurons are located in three anatomical groups in the ventral midbrain, namely the ventral tegmental area (VTA, A10), the substantia nigra pars compacta (SNc, A9) and the retrorubral field (RRF, A8; Watabe-Uchida et al. 2012; Beier et al. 2015; Morales and Margolis 2017; Bjorklund and Dunnett 2007; Pearson et al. 1983; Arsenault et al. 1988). In primates, the lateral part of the VTA, the parabrachial pigmented nucleus (PBP, A10), extends from the midline VTA (A10), to stretch dorsally across the mediolateral extent of the ventral midbrain, and transitions caudally into the RRF (A8; Fudge et al. 2017; Halliday and Tork 1986). The PBP and A8 are thus disproportionately expanded territories in nonhuman and human primate, compared to other regions (Halliday and Tork 1986; Fu et al. 2016; McRitchie et al. 1996; Francois et al. 1999). Each of the DA subregions is composed of cells with diverse phenotypic characteristics, and are differentially connected with various brain areas correlating with their anatomic position in the ventral midbrain (Arias-Carrion et al. 2010; Lammel et al. 2011; Lammel et al. 2014; Farassat et al. 2019; Haber et al. 2000). As one example of cellular heterogeneity across subregions, we recently found a broad range of DA:GABA cell ratios in different subregions in monkey, suggesting differences in intrinsic control of DA firing (Kelly et al. 2022).

The DA neurons receive excitatory, inhibitory, and modulatory input from many sources (Lewis and Sesack 1997; Morales and Margolis 2017; Watabe-Uchida et al. 2012). Corticotropin-releasing factor (CRF), a canonical 'stress' peptide, is found in some of these afferent sources, and is a neuromodulator of DA cell firing, although the precise mechanism is not understood (Kalivas et al. 1987; Ungless et al. 2003; Wanat et al. 2008; Wanat et al. 2013; Grieder et al. 2014; Gallagher et al. 2008; Orozco-Cabal et al. 2006). Like all neuropeptides, CRF is always expressed with a primary transmitter. CRF is synthesized, ribosomally, packaged and modified in dense core vesicles (dcv), then transported down the axon shaft to the terminal (Hokfelt et al. 2003). At the terminal, punctate CRF-immunoreactivity is found surrounding small clear vesicles that typically sequester ‘fast’ primary transmitters, and in dense core vesicles (dcv) (Van Bockstaele et al. 1998, 1996). CRF-containing terminals may have asymmetric (putative excitatory) or symmetric (putative inhibitory) synaptic specializations, dictated by the primary transmitter with which it is packaged (Henckens et al. 2016). Potential sources of CRF to the various DA subregions are therefore diverse, but have not been well studied (reviewed in, Kelly and Fudge 2018). A large known source of CRF to the rodent and primate ventral midbrain DA system in general arises from the central extended amygdala (Fudge et al. 2017; Dabrowska et al. 2016; Rodaros et al. 2007).

In rodents, the ventral tegmental area (VTA) is seen as an important effector site for CRF action, due to its outputs to the ventral striatum and medial prefrontal cortex in that species (reviewed in, Morales and Margolis 2017; Bjorklund and Dunnett 2007). In primates, evolutionary extension of the ventral midbrain results in differential expansions of VTA subnuclei (A10), and also the A9, and A8 regions. With this expansion, the pattern of efferent and afferent circuits through these areas also shifts (reviewed in, Bjorklund and Dunnett 2007; Cho and Fudge 2010; Fudge et al. 2017). It is now recognized that the broad expanse of the primate DA system is physiologically heterogeneous across the mediolateral axis with respect to both intrinsic firing, and coding properties. DA neurons in the 'midline' VTA nuclei code reward prediction errors (Mirenowicz and Schultz 1994; Tobler et al. 2003), while more dorsolaterally placed DA neurons in the vicinity of the PBP and A8, are 'salience' coding, i.e. respond to potentially salient stimuli regardless of value (Matsumoto and Hikosaka 2009; Bromberg-Martin et al. 2010). These differences suggest that CRF terminals in the 'midline' VTA versus PBP/A8 subpopulations may have differing functional roles, since each region forms specific circuits (Bromberg-Martin et al. 2010; Fudge et al. 2017; Haber et al. 2000). The organization of CRF terminals in DA subpopulations outside of the classic VTA -- ie., in regions associated with 'salience' coding of noxious and novel stimuli--is of interest.

In this study, we investigated the interactions of CRF-positive synaptic terminals in the PBP and A8 subregions in young male and female macaques. Using dual immuno-peroxidase reactivity for electron microscopy (EM), we quantified ultrastructural synaptic contacts of CRF-reactive fibers onto either tyrosine hydroxylase (TH) positive and TH-negative neurons in the PBP and A8 subregions.

## MATERIAL AND METHODS

### Animals

All experiments were conducted in accordance with National Institute of Health guidelines (NIH Publications No. 80-23). Experimental design and technique were aimed at minimizing animal use and suffering and were reviewed by the University of Rochester Committee on Animal Research. Animals were all socially housed by sex. To conserve animals, we used perfused tissue from 3 young male and 3 young female monkeys (*Macaque fascicularis*) that had received tracer injections in other brain regions as part of other studies (**Table 1**). The three female animals were also part of a larger cohort examined for longitudinal hormone studies to ascertain general stress and neurodevelopment state in same-age animals pair-housed in our facility.

**Table 1.** Experimental Cases

| Case | Weight<br>(kg) | Age<br>(years) | Sex |
| --- | --- | --- | --- |
| J32 | 5.4 | 3.3 | M |
| J38 | 3.5 | 3.3 | M |
| J41 | 4.1 | 4.5 | M |
| J55 | 3.5 | 2.9 | F |
| J58 | 4.0 | 2.9 | F |
| J59 | 3.1 | 3.1 | F |

### Histology

All animals were sacrificed under deep anesthesia (pentobarbital) by intracardiac perfusion, first with 0.9% sterile saline and then 6L of 4% paraformaldehyde (PFA) in a solution of sucrose and phosphate buffer (PB, pH 7.4). Brains were harvested and placed in 4% PFA overnight, and then sunk in increasing gradients of sucrose solution (10%, 20% and 30%). The entire brain was coronally sectioned on a freezing sliding microtome at 40 um and all sections were saved in serial wells (24 serial ‘compartments’; 1:24) in a cold cryoprotectant solution containing 30% sucrose and 30% ethylene glycol in PB at −20°.

### Antibody Characterization

The antibodies used in this study are detailed in **Table 2**. Anti-CRF (rabbit, T-4037, Peninsula) shows specific localization of CRF in axon terminals (Tagliaferro and Morales 2008; Kelly and Fudge 2018; Fudge et al. 2017; Yuan et al. 2019) and has been thoroughly tested using pre-absorption assays (JCN antibody database ID #AB-518252). In primate, we have previously shown specific immunoreactivity (IR) in CRF positive cells and fibers in the macaque brain (Kelly and Fudge 2018; Fudge et al. 2017), with similar findings as reports using other CRF antibodies (Cha and Foote; Bassett and Foote). Anti-tyrosine hydroxylase reactivity (TH, mouse, MAB518, Millipore, clone LNC1) specificity is well-documented in many species (JCN antibody database ID # AB-2201528)(Cho and Fudge 2010), and our results are in agreement with previously reported distribution and cytoarchitecture of TH-IR cells in monkey (Arsenault et al. 1988). TH-IR is an accepted marker of DA cells in the ventral midbrain, as enzymes involved in the synthesis of norepinephrine or epinephrine are not detectable in this region (Pearson et al. 1983; Gaspar et al. 1983). The pattern of calbindin-D28K (CaBP) expression is also identical to published reports for primate A10 and A8 dopamine neurons, and has been authenticated previously(Cote et al. 1991; McRitchie et al. 1996; Gaspar et al. 1993) (JCN antibody database ID #AB-476894).

**Table 2.** Antibodies Used

| Antigen | Immunogen | Source | Characterization Reference | Working Dilution |
| --- | --- | --- | --- | --- |
| Corticotropin Releasing Factor (CRF)<br>Host: Rabbit | Polyclonal antibody collected from rabbits immunized with a synthetic peptide as the immunogen. Species cross reactivity: human, mouse, rat. | Peninsula Laboratories, Inc. (Bachem Group)<br>T-4037 | *Journal of Comparative Neurology* antibody database ID #**AB-518252** | 1:1,000 |
| Tyrosine hydroxylase (TH)<br>Host: Mouse | Monoclonal antibody, clone LNC1. Recognizes an epitope on the outside of the regulatory N-terminus. Species cross-reactivity: human, mouse, rat, chicken, frog, monkey, and vole. | Millipore,<br>MAB318 | *Journal of Comparative Neurology* antibody database ID #**AB-2201528** | 1:10,000 |
| Calbindin (CaBP)<br>Host: Mouse | Monoclonal antibody, clone CB-955. Highly conserved 28kD calcium binding protein with broad tissue distribution. Species cross-reactivity: rat, bovine, guinea pig, human, mouse, canine, sheep, rabbit, feline, goat, pig. | Sigma, C9848 | *Journal of Comparative Neurology* antibody database ID #**AB-476894** | 1:10,000 |

### Single-label immunohistochemistry

For region of interest (ROI) localization in EM studies, near adjacent sections were immunostained for CRF, CaBP and TH (**Figure 1**). This approach allowed us to sample from areas with known dense termination of CRF positive fibers specifically in the PBP or A8, confirmed by matching microscopic anatomical landmarks (blood vessels patterns, *) for accurate excision (trapazoid region for orientation) of PBP and A8 blocks for EM samples. Near adjacent compartments of tissue were singly immunoreacted for anti-Calbindin D28k (CaBP,a marker of the A10 and A8 DA subpopulation 1:10K, Sigma #C9848, mouse,Haber and Fudge 1997), anti-CRF (1:1K, Penisula #T-4037, rabbit) and anti-TH, 1:10K, Millipore #MAB318, mouse). Briefly, sections were rinsed thoroughly in 0.1M phosphate buffer (PB, pH 7.4) containing 0.3% Triton-X (TX) followed by an endogenous peroxidase step (10% methanol, 3% H_2_O_2_ in 0.1M PB). Sections were then rinsed thoroughly in PB-TX and blocked with 10% normal goat serum in PB- TX (NGS-PB-TX; primary antisera). Following rinses in PB-TX, tissues were incubated in primary antisera for ∼96 hours at 4°C. Tissues were then rinsed with PB-TX, blocked with 10% NGS-PB-TX, incubated in the appropriate biotinylated secondary antibody (goat anti-rabbit, #BA-100; goat anti-mouse, Vector Labs #BA-9200;Vector Laboratories, Burlington, ON Canada) and then incubated with avidin-biotin complex (Vectastain ABC kit, PK-6100 (ABC Elite, for CaBP and TH) or PK-4000 (Standard ABC kit, for CRF), Vector Laboratories, Burlington, ON Canada). After rinsing, sections were visualized with 3, 3’- diaminobenzidine (DAB, 0.05mg/ml, 0.3% H_2_O_2_, in 0.1M Tris-buffer), and rinsed thoroughly prior to mounting. Sections were mounted out of mounting solution (0.5% gelatin in 20% ETOH in double distilled water) onto subbed slides, dried for 3 days, and cover slipped with DPX mounting media (Electron Microscopy Sciences).

**Figure 1.**
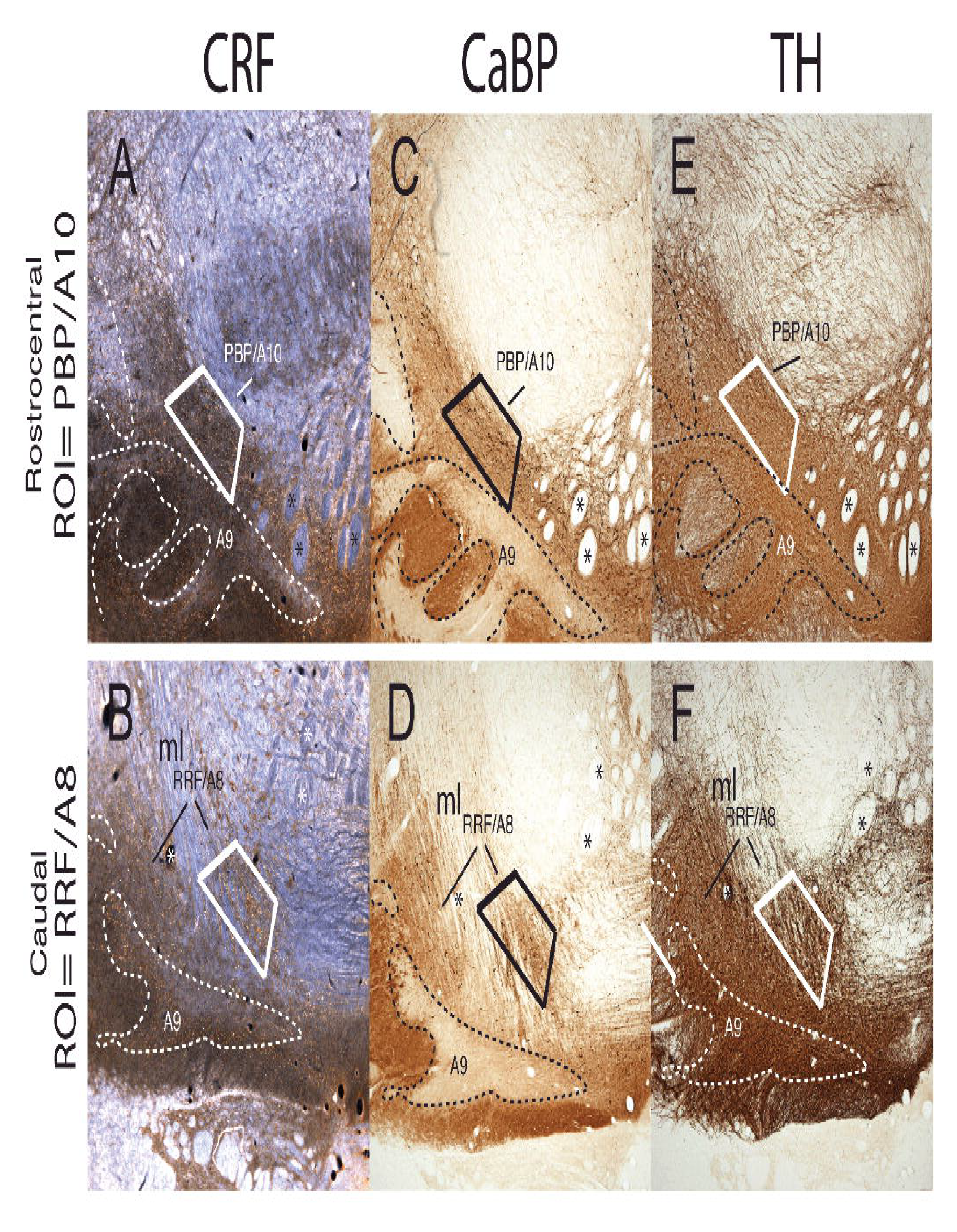
Localization of the PBP and A8 for EM. Near adjacent compartments immuno-labeled for CRF, CaBP and TH to identify and block regions of interest (ROIs) for immuno-EM processing. Rostral adjacent compartments showing the PBP (**A, C, E**) and caudal adjacent compartments containing A8 (**B, D, F**) with CRF-, CaBP-, and TH-IR, respectively. Fiducial markers such as fiber bundles (e.g. fascicles of the 3^rd^ nerve(asterisks) and medial lemniscus (ml)) and blood vessels were used to localize the region of interest in EM section.

### Double-immunoperoxidase and immunogold-reactivity for CRF and TH (EM)

Brain sections adjacent to those used for ROI identification (**Figure 1**) were dually stained for CRF-IR and TH-IR. Briefly, sections were washed thoroughly in freshly-made filtered 0.05M phosphate buffered saline (PBS), subjected to 0.1% sodium borohydride in 0.1M filtered 0.1M PB to block active aldehydes (Craig 1974), and blocked in a solution containing 3% NGS and 1% bovine serum albumin (BSA). Sections were incubated for 72-hours in the same blocking solution with rabbit anti-CRF (Peninsula,T-4037, 1:1K) and mouse anti-TH (Millipore, MAB318, 1:5K) at 4°C on a shaker followed by 24-hours at room temperature (RT). Sections were then washed in filtered 0.05M PBS and incubated for 1 hour in goat anti-rabbit IgG conjugated to biotin (1:200, Jackson ImmunoResearch, BA-1000), followed by ABC (per product instructions; Elite ABC Kit, Vector Labs, PK-6100). CRF immunoreactivity was visualized with 3,3’- diaminobenzidine (0.5mg/ml; DAB) and hydrogen peroxide (0.03%) in 0.1M PB. Sections were rinsed thoroughly in filtered 0.05M PBS, followed by a second pre-incubation in a washing buffer solution containing 3% NGS, 0.8% BSA, 0.1% cold water fish gelatin in 0.1M PBS.

Sections were then incubated for 24 hours in goat-anti-mouse immunogold (Nanogold, N24915, Nanoprobes Inc., Yaphank, NY) in washing buffer at RT on a shaker. Following a thorough wash, tissue was incubated in 2.5% glutaraldehyde for 10 minutes at RT. Sections were rinsed thoroughly in 0.1M PBS, followed by a series of washes in Enhancement Conditioning Solution (1X concentration per package instructions, Electron Microscopy Science (EMS), Cat 25830, Hatfield PA). Sections were silver enhanced using the Aurion R-Gent SE-EM Kit (EMS, Cat #25520-90). Following additional post-fixation in 1% osmium tetroxide (EMS, Cat #19150), sections were dehydrated in ascending concentrations of ethanol (50%, 70%, 80%, 95%, 100% x2). Contrast was enhanced using 1% uranyl acetate in 70% ETOH within the dehydration series (EMS, Cat 22400). Sections were then treated with propylene oxide, impregnated in resin (Embed 812, Araldite 502, DDSA, DMP-30 (all EMS)) overnight at RT, mounted between ACLAR embedding films (EMS, Cat# 50425) and cured at 55°C for 72-hours. Using immunolabeled adjacent sections to carefully identify them (**Figure 1**), the PBP and A8 regions were excised in a trapezoid shape (to record anatomical orientation) from the embedding films and re-embedded at the tip of resin blocks. Ultrathin sections (60-80 nm; evidenced by the sections silver sheen) were cut with an ultramicrotome (Reichert Ultracut E) and collected in sequence on bare square-mesh nickel grids (EMS, G200-Ni).

### EM Imaging and Data Analysis

Images were captured on a Hitachi 7650 Transmission Electron Microscope using a Gatan 11-megapixel Erlangshen digital camera and Digital Micrograph software. TIFF images were later exported into Adobe Photoshop and Adobe Illustrator (v2022) and adjusted for brightness and contrast in preparation for analysis. Starting at the upper left-hand corner of each grid opening (selected at least 10 microns from the tissue: resin interface), tissue was first visualized at 800x, then the area was subsequently surveyed under higher power (12,000-40,000x) to discern positive CRF-positive elements (**Figure 2A, B, C**). Samples with poor staining or tissue preservation were excluded from further analysis. Eighty to one-hundred electron micrographs containing CRF immunoreactive elements were arbitrarily taken at a final magnification of 30,000-40,000x, for quantification (see Quantification, below). These photomicrographs, at this magnification, correspond to a total surface of ∼ 1,000 μm^2^ of neuropil per region per animal (as in, Kelly et al. 2014; Tremblay et al. 2007; Bouvier et al. 2008). In addition, 70,000 magnification micrographs were collected in many instances to confirm findings (**Figure 2D**), and create presentation images. Gold particle processing was titrated to diffusely label TH-positive structures, enabling clear views with respect to synaptic structures. A lower magnified image of the structure was first referenced to ensure an accurate classification of TH-positive elements (**e.g. Figure 2**) prior to taking final photomicrographs at higher magnification, i.e. 30, 000-40,000x, and 70,000x magnification images for some ROIs.

**Figure 2.**
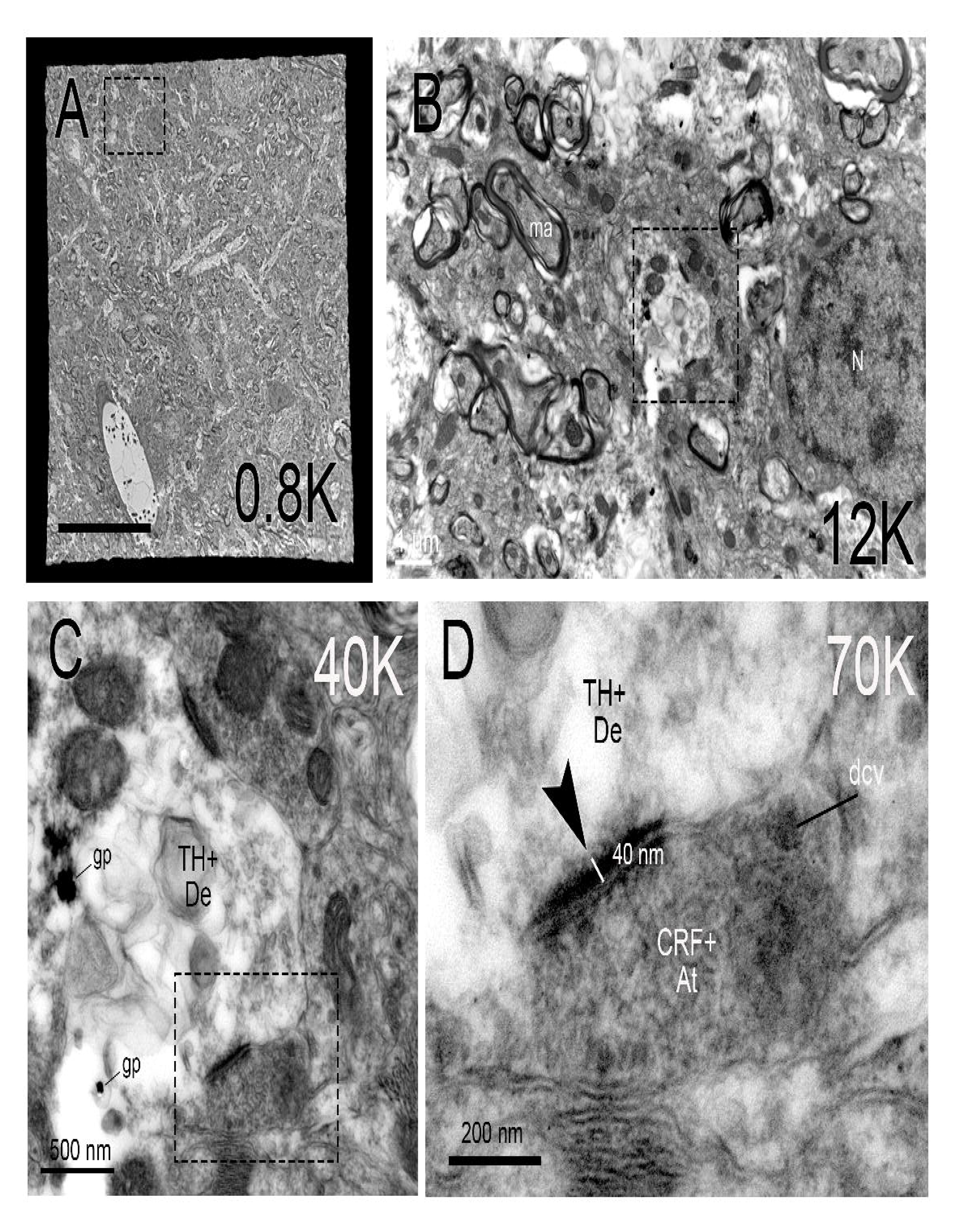
EM micrograph acquisition. (**A**) Example of a single electron microscopy grid opening at 0.8 K magnification. Scale bar= 20 um (**B**) 12K magnification of boxed region in (A) showing visualization of synaptic vesicle-rich presynaptic terminals and electron-lucent dendritic structures (with and without gold particle deposits). Scale bar= 1 um (**C**) Under 40K magnification, boxed region in (B) shows synaptic specialization and a dense core vesicle in the axon terminal. At this magnification, pre-and postsynaptic membranes are discernible, CRF+ immunoreactive accumulation is seen around synaptic vesicles, and some dcvs are visible. Two gold particles identify the post-synaptic dendrite as TH-positive (TH+De) Scale bar= 500 nm (**D**) 70K magnification of boxed region in (C) to confirm structural identificaiton. A CRF+ immunoreactive presynaptic terminal (CRF+At) abuts the TH-positive dendrite, identified in C. A dense core vesicle is apparent at the outer edge of the asymmetric synapse (black arrow), in addition punctate reaction product. The asymmetric synapse is classified by the presence of a 40nm post-synaptic density, and little to no density on the presynaptic side. Scale bar=200 nm. *Abbreviations:* CRF+At, CRF-positive axon terminal; dcv, dense core vesicle; gp, gold particle; ma, myelinated axon; N, nucleus; TH+DE, TH- positive dendrite.

Specific cellular components were identified using a series of criteria previously defined in single-ultrathin sections (Peters et al. 1991; Kelly et al. 2014). In instances when an immunoreactive density obscured some of the intracellular organelles and defining characteristics, we relied on the visible defining features, such as contours made by the profiles and ultrastructural relationships with neighboring elements to aid in our classification. The following classifications were applied to characterize labeled axons, axon terminals, and dendritic elements throughout the neuropil, which were the main structures of interest.

*Axon terminals, axon shafts*- Axonal terminals were distinguished from other subcellular profiles based primarily on the presence of small synaptic vesicles, resulting in an electron-dense appearance. Axon terminals generally contained mitochondria. Axon terminals were further characterized based on synaptic contact with neighboring elements. In contrast to terminals, axon shafts appeared round when viewed in the transverse plane (i.e. axon bundles) and elongated when cut longitudinally. In CRF labeled axon terminals, punctate DAB accumulation was evident around small, clear, unlabeled synaptic vesicles, similar to previous reports, e.g. (Van Bockstaele et al. 1996; Tagliaferro and Morales 2008). CRF- immunoreactivity was also seen in dense core vesicles, which were localized among clusters of small vesicles, and often bordering or close to active zone (Persoon et al. 2018). Dense core vesicles were larger than small clear vesicles (approximately 100-250 nm, versus 40 nm for small vesicles), consistent with the literature (Peters et al. 1991). In our material, dense core vesicles with CRF-positive and CRF- negative cores were seen. Labeled axon shafts showed similar patterns of immunoreactive punctae along with occasional dense core vesicles.

*Dendritic shafts*- Dendritic shafts cut longitudinally were recognized by their irregular contours, elongated mitochondria in parallel with their central axis, frequent protuberances (spines, filopodia, small branches), and synaptic contacts with axon terminals. When cut transversally, dendritic shafts were identified by their rounded morphology, frequent occurrence of mitochondria and microtubules, and were distinguished from unmyelinated axons by their larger diameter. Dendrites appear more electron-lucent than axon shafts. For TH-positive dendrite identification, the dendrite was considered TH-positive if gold particles were present in the entire structure at lower power. The structure was noted as TH-positive, and imaged at higher power.

*Dendritic Spines*- Dendritic spines cut longitudinally often protruded from dendritic shafts, displayed rounded morphologies and were free of mitochondria. Spines were characterized primarily by the presence of electron-dense accumulations (postsynaptic densities) at synaptic contact sites with axon terminals.

*Astrocytes*- Protoplastic astrocytes were recognized as electron-lucent structures seen to encase and wrap around other neuropil structures. As a result, astrocytes maintained irregular and angular shapes, distinguishing them from other neuronal profiles having a characteristic rounded shape.

### Quantitative Analysis

All electron micrographs were analyzed by one investigator (EAK) blinded to animal sex. Only CRF- positive terminals with synaptic profiles were counted. ‘Synapses’ were classified as CRF-IR axon terminals that made either a symmetrical or asymmetrical synaptic profile onto a post-synaptic element (usually a dendrite). A synaptic terminal was considered CRF-positive (CRF-At+) if it contained either punctate DAB deposits or immunoreactive dense core vesicles, or both. We did not count CRF terminals making ‘appositions’, where the CRF-labeled presynaptic membrane was apposed to the postsynaptic membrane, but there was no apparent synaptic specialization. All synapses were identified by thickened parallel membranes with a widened cleft. Symmetric synapses had a thin post-synaptic densities characteristic of inhibitory synapses. Asymmetric synapses had thickened post-synaptic densities (PSDs), generally greater or equal to 40nm (Dosemeci et al. 2016; Root et al. 2018; Peters et al. 1991) (**Figure 2C, D**).

In each micrograph, the number of CRF-labeled axon terminals was determined. Then, synapse contact type for each terminal was assessed based on the criteria above. The total number of asymmetric and symmetric synapses were calculated for each case. We then determined whether CRF positive terminals made a synaptic contact with TH-positive (TH+De) or TH-negative (TH-De) dendritic profiles based on the presence or absence of immunogold labeling. Because gold labeling was titrated so as to not obstruct classification of synaptic contacts, visualization of large post-synaptic densities (PSDs) that distinguish asymmetric (excitatory, black arrow heads) synapses, from the thin PSDs that make up symmetric (inhibitory, white arrow heads) synapses, was possible (**Figure 2**).

Our analyses focused on: (1) the total number of CRF-positive axon terminals with synaptic contacts on TH-positive versus TH-negative dendritic profiles, (2) the proportion of asymmetric vs symmetric synapses across TH-positive and TH-negative dendritic profiles in the PBP versus A8, and (3) the proportion of asymmetric and symmetric synapses onto either TH-positive or TH-negative dendrites in each region. Proportions were determined (1) dividing the number of asymmetric or symmetric synapses by the total number of synapses and (2) by dividing the total number of each synapse type by the total number of each dendritic profile (TH-positive versus TH-negative).

### Hormone Assays

Serum hormonal specimens were collected and assayed on 7 animals (n= 3 males, n=4 females) kept in our facility in 2020. Three of these animals (female) were included in the EM study. The purpose was to assess the long-term stability of pubertal and stress measures in same-age male and female animals under usual conditions (pair housing) maintained in our facility. Specimens were collected at three time points over approximately 6 months: 1) when animals were released from quarantine (60 days after arrival to the facility), 2) before tract-tracer surgery (approximately 30-90 days after release from quarantine) and 1. 3) on the morning of sacrifice (approximately 2 weeks after surgery). All samples were collected at the University of Rochester and shipped on dry ice for hormone analysis off-site. At least 2 mL of serum (from 4mL whole blood) was collected using either an IV catheter or butterfly needle/syringe into red topped tubes. Blood was then centrifuged, and serum pipetted off and stored at −20. Assays were run at the Wisconsin National Primate Research Center, under the supervision of Dr. Amita Kapoor. The following references provide a detailed report of the techniques used for these analyses: Multi-steroid Assays from Serum: (Kenealy et al. 2016) and radioimmunoassay for luteinizing hormone (Garcia et al. 2018).

### Statistics

Analysis was performed with Prism 9 software (GraphPad Software) and R software. All values reported in the text are mean ± standard error of the mean (SEM). For all statistical tests, significance was set to p<0.05. Analysis was conducted using a generalized linear model (GLM) with a single dependent variable and two or three independent categorical variables and corrected Tukey’s multiple comparisons post-hoc contrasts (comparing each cell with every other cell). For dependent variables measuring synaptic contact counts, a Poisson family with a log-link was used. For dependent variables measuring proportions of synaptic contacts, a binomial family with a logit link was used. Within and across independent variable comparisons are denoted with significance bars in each figure panel accordingly. In detail, we made the following comparisons: Figure 6A: The total number of CRF+ synaptic contacts (dependent variable; DV) within the TH-positive and TH-negative profile populations (independent variable 1; IA-1) and DA subregions (PBP or A8; independent variable 2; IA-2). Figure 6B: The proportion of CRF+ synaptic contacts (DV) within the asymmetric synapse and symmetric synapse populations (IA-1) and DA subregions (PBP or A8; IA-2). Figure 6C: The proportion of CRF+ synaptic contacts (DV) within the TH- positive and TH-negative profile populations (IA-1) across the asymmetric synapse and symmetric synapse populations (IA-2) in PBP/A10. Figure 6D: The proportion of CRF+ synaptic contacts (DV) within the TH- positive and TH-negative profile populations (IA-1) across the asymmetric synapse and symmetric synapse populations (IA-2) in RRF/A8. Figure 7A: The total number of CRF+ synaptic contacts (DV) within the TH- positive and TH-negative profile populations (IA-1) in both male and female populations (IA-2) in PBP/A10. Figure 7B: The total number of CRF+ synaptic contacts (DV) within the TH-positive and TH-negative profile populations (IA-1) in both male and female populations (IA-2) in RRF/A8. Figure 7C: The proportion of CRF-positive synaptic contacts (DV) across the asymmetric synapse and symmetric synapse populations (IA-1), gender (Male vs Female; IA-2), and TH-positive and TH-negative profiles (independent variable 3; IA-3) in PBP/A10. Figure 7D: The proportion of CRF-positive synaptic contacts (DV) across the asymmetric synapse and symmetric synapse populations (IA-1), gender (Male vs Female; IA-2) and TH-positive and TH-negative profiles (IA-3) in RRF/A8.

## RESULTS

### CRF-positive axon terminals in the ventral midbrain subregions: light microscopy

The A10 neurons in the nonhuman primate extend the entire rostrocaudal extent of the midbrain (approximately 6 mm), and include the following subnuclei: the rostral linear nucleus (RLi), the caudolinear nucleus (CLi), the intrafascicular nucleus, the ventral tegmental nucleus (VTA), the paranigral nucleus, and the parabrachial pigmented nucleus (PBP, Halliday and Tork 1986; McRitchie et al. 1995). In human and in monkey, the PBP is by far the largest sub-nucleus of the A10 (Halliday and Tork 1986; Olszewski and Baxter 2014), and sweeps dorsolaterally over the entire A9 region, to merge caudally with the A8 group. We therefore grouped all individual VTA subnuclei medial to the PBP as the ‘midline VTA’ group and designated the PBP as a separate A10 subnucleus. PBP TH-positive neurons are frequently larger than other A10 neurons, and oriented horizontally with long proximal dendrites in the mediolateral plane. CaBP cellular immunoreactivity marks the A10 and A8 DA subregions across species (Gaspar et al. 1993; Haber and Fudge 1997; Lavoie and Parent 1991; McRitchie et al. 1996), with noticeable absence in the SNc (A9) DA subregion (**Figure 3A, rostrocentral, 3D, caudal**).

**Figure 3.**
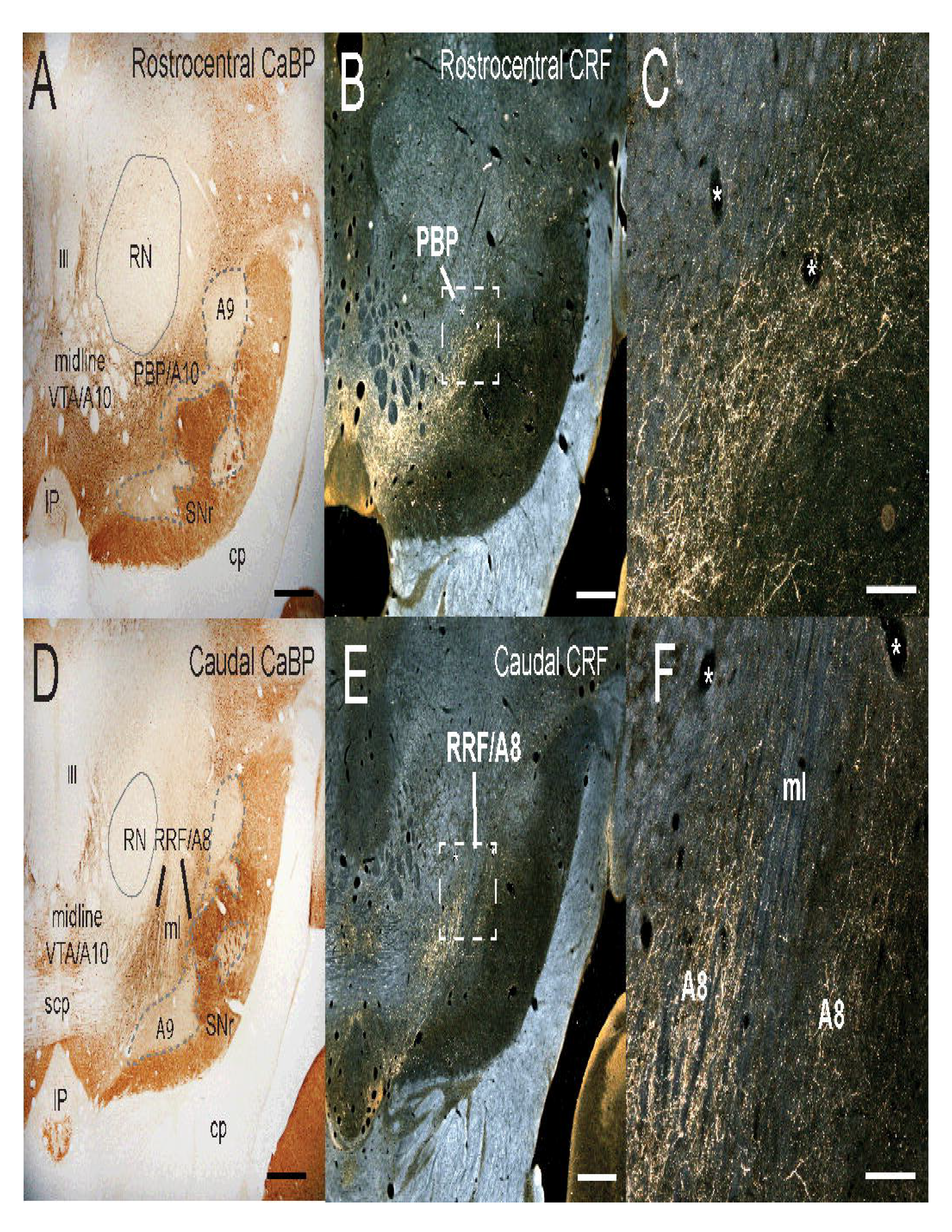
Localization of CRF immunoreactivity in ventral midbrain. **(A)** Low magnification brightfield micrograph of the ventral midbrain rostrocentral level. CaBP-IR, a marker of the A10 and A8 neurons. CaBP-positive A10 neurons contrast with CaBP-negative SNc/A9 neurons, outlined with dotted line. The substantia nigra, pars reticulata (SNr) has dense CaBP-IR fibers. **(B)** CRF-IR in fibers in adjacent section, visualized with darkfield microscopy. **(C)** A higher magnification of boxed region in B showing patches of thin beaded CRF- positive fibers in a section of the PBP. **(D)** Low magnification brightfield micrograph of CaBP-IR in the caudal ventral midbrain. CaBP-IR neurons are found in the A10 and RRF/A8, and absent in SNc/A9 subregion. **(E)** CRF-IR in a neighboring section to panel D, seen under dark-field microscopy. **(F)** CRF- labeled fibers course through the RRF/A8, which is bi-sected by the medial lemniscus (ml) seen under higher magnification (taken from boxed area in E). Scale bar in A,B,D,E= 1 mm; C,F= 250 um. *Abbreviations:* III, Third nerve; CRF, corticotropin releasing factor; CaBP, calbindin-28kD; IP, interpeduncular nucleus; cp, cerebral peduncle; scp, superior cerebellar peduncle; ml, medial lemniscus; SNr, substantia nigra reticulata; PBP, parabrachial pigmented nucleus; RN, red nucleus; VTA, ventral tegmental area.

Comparing the distribution of CRF-positive terminals with adjacent CaBP-labeled sections at the light microscopic level, robust CRF innervation was found predominately rostrocentrally over the PBP, and at caudal levels over the RRF/A8 region (Figure 1B-C [rostrocentral], Figure 1E-F [caudal]; Kelly and Fudge 2018). There were few labeled CRF-positive fibers extending into the CaBP-negative A9 region. As expected, CRF-positive axonal terminals were also found in the midline VTA nuclei, where they were of a higher density at rostrocentral compared to caudal levels (**Figure 3B; 3E**). In both the PBP and A8, dense patches of CRF-labeled fibers consisted of very thin, highly varicose fibers, as well as some thicker, beaded 'fibers en passant' (PBP, **Figure 3C;** RRF **/**A8, **Figure 3F**).

### Characteristics of CRF-labeled axon terminals and TH-positive dendrites: electron microscopy

At the EM level, CRF immunoreactivity was predominantly found in axon shafts and terminals (**Figure 4**) and only rarely in dendrites (2 instances, not shown). Depending on the plane of sectioning, CRF immunoreactive axon terminals contained punctate immuno-peroxidase deposits (See **Figure 4A**) interspersed among small, clear synaptic vesicles, and in some, CRF-positive large dense core vesicles (dcvs, approximately 80-200 nm)(**Figure 4A, D**). Unlabeled dense core vesicles were also found within CRF-positive and CRF-negative terminals, consistent with the presence of other neuropeptides in afferent terminals, and/or recently released CRF (e.g. **Figure 5B,** udcv) (VanBockstaele et al. 1996; Maley 1990; Nusbaum et al. 2017). CRF positive axon terminals made asymmetric and symmetric synaptic contacts, denoting excitatory and inhibitory synapses, respectively (e.g **Fig. 4A,C,D**). Both types of synaptic profiles were found on TH-positive and TH-negative dendrites. CRF-labeled axon terminals were also frequently enveloped by an astrocytic process (asterisks in **Figure 4B,D**) as documented by others (VanBockstaele et al. 1996).

TH immunoreactivity, visualized using gold/silver enhancement, was found in dendritic structures and cell bodies (**Figure 5A-C**). The majority of TH immunoreactivity was in medium to large dendritic structures (1- 2μm diameter). Single TH-positive dendrites often displayed multiple synaptic contacts, both symmetric and asymmetric in nature. Non-TH immunoreactive dendrites were identified as having similar ultrastructural features, but without gold particles. Small offshoots of TH-immunoreactive structures (such as spines and thin dendritic elements) may have been left unlabeled due to our gold particle titration, and therefore underrepresented in our analysis.

**Figure 4.**
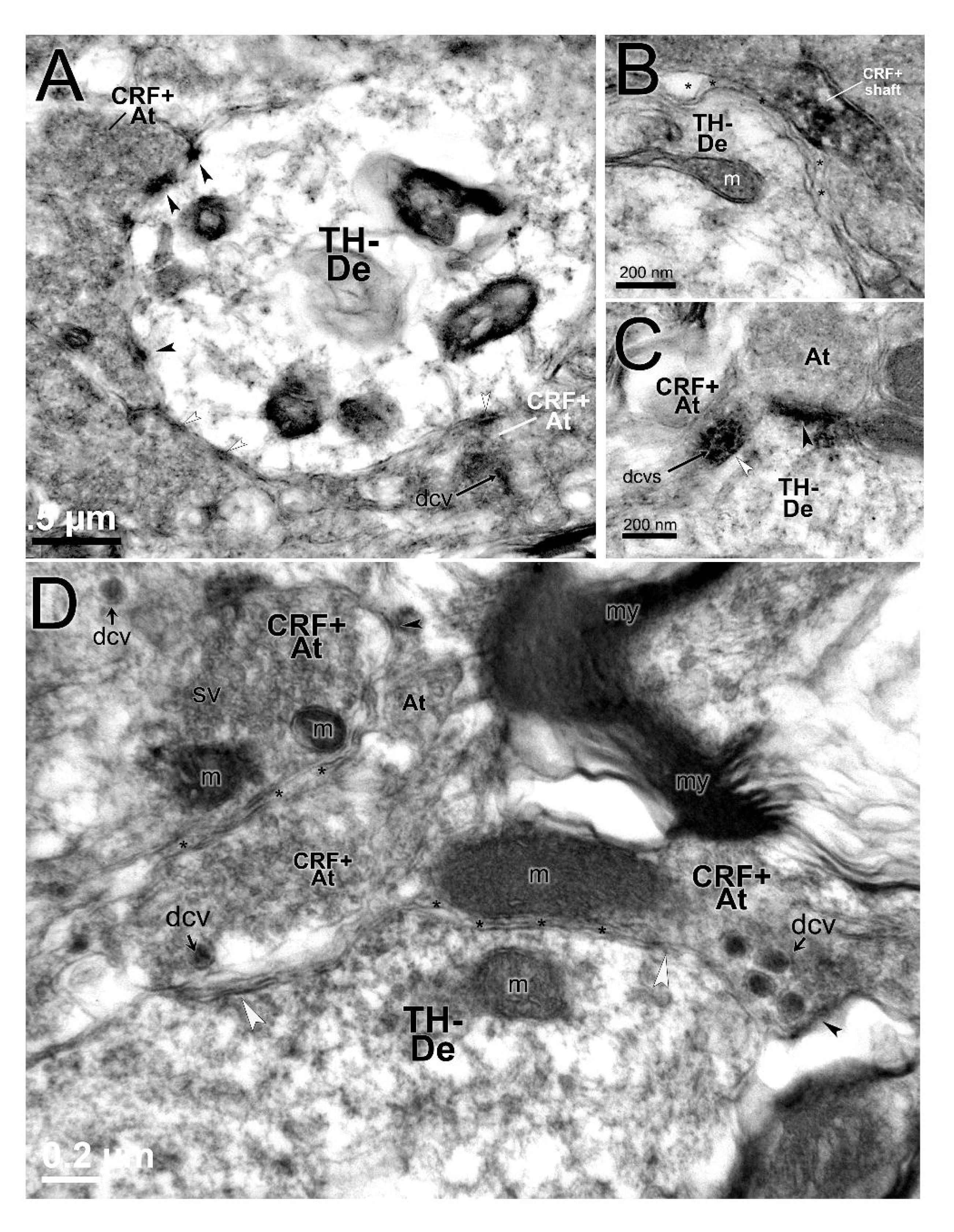
CRF-IR axon terminals. **(A)** Several CRF-IR axon terminals (CRF+At) with synaptic specializations on a TH- negative, electron-lucent dendrite. CRF-IR is seen as irregular peroxidase labeling surrounding clear vesicles throughout the presynaptic space, and in a dense core vesicle (dcv, arrow) which lies away from the synaptic junction. Scale bar= 500nm (**B**) A CRF-labeled axonal shaft (CRF + shaft), enveloped by a clear astrocytic process (asterisks) neighboring a TH-negative dendrite. Scale bar= 200 nm. **(C)** CRF+ At, with CRF+ dcvs, making a symmetric synapse (white arrowhead) with a TH-negative dendrite. Adjacent is an At (CRF-negative) making an asymmetric contact (black arrowhead) with the same dendrite. Scale bar=200 nm. (**D**) CRF-positive axon terminals (CRF-At+); several are abutting a TH-negative dendrite (TH- De). A cluster of darkly labeled dense core vesicles (dcvs) are evident in the CRF-At+ structure on the right. Asterisks show astrocytic process interposed between the terminal and dendrite. Symmetric (white arrowheads) and asymmetric (black arrowheads) synaptic profiles are indicated. Scale bar=200 nm. *Abbreviations:* At, axon terminal (unlabeled); CRF+At, CRF-positive axon terminal; dcv, dense core vesicle; m, mitochondria; ma, myelinated axon; sv, small vesicles; TH+De, TH-positive dendrite; TH- dendrite, TH-negative dendrite.

**Figure 5.**
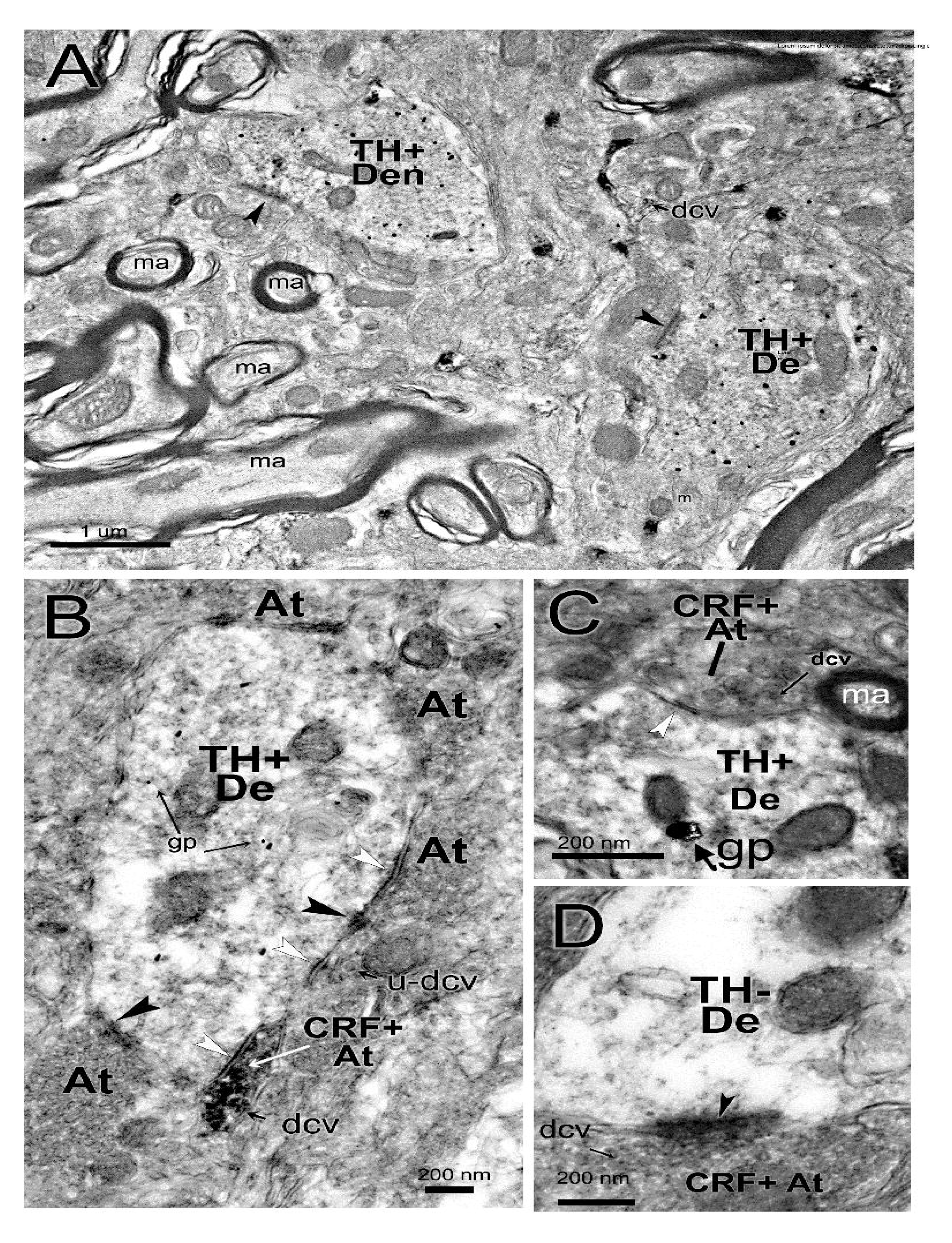
CRF-positive terminals onto TH-positive and TH-negative dendrites. **(A)** Lower magnification (30,0000X) electron micrograph detailing the specificity of TH-positive gold particles in several dendritic structures (TH+De). Gold particles are diffusely distributed throughout the cytoplasm of electron-lucent dendrites, allowing characterization of synaptic contacts by neighboring presynaptic terminals at higher power. Black arrowheads, asymmetric synapses; small arrow indicates appearance of a dense core vesicle (dcv) at low power. Scale bar= 1 micron. **(B)** A CRF-positive axon terminal (CRF+At) with dcv and punctate immunoreactive immunoreactivity making a symmetric contact (white arrowhead) with a TH-positive dendrite (TH+De). Several gold particles (gp) are seen in the dendrite. Other unlabeled axon terminals make contact with the same TH+De (black arrowheads, asymmetric synapse; white arrowheads, symmetric synapse). An unlabeled dense core vesicle (u-dcv) is also indicated. Scale bar= 200 nm. (**C**) A CRF-positive axon terminal (CRF+At) with several CRF-positive dcvs (arrow) making a symmetric contact with a TH-positive dendrite. Scale bar-200 nm. (**D**) A large CRF-positive axon terminal (CRF+At) with dcv making an asymmetric (excitatory) synaptic contact onto a TH-negative dendrite (black arrowhead). Scale bar-200 nm. *Abbreviations:* At, axon terminal (unlabeled); CRF+At, CRF-positive axon synaptic terminal; dcv, dense core vesicle; gp, gold particle; m, mitochondria; ma, myelinated axon; TH+De, TH-positive dendrite; TH-De, TH-negative dendrite, u-dcv, unlabeled dense core vesicle.

### CRF-positive terminals synapse mainly onto TH-negative dendrites predominate in both PBP and A8

We first quantified CRF-positive terminals with synapses on each dendritic type in the PBP and A8 (**Figure 6A**). The majority of CRF-positive terminals with recognizable synapses were predominantly onto TH- negative dendrites in both regions: the PBP (89%; 917 synapses: 1031 total synapses; black bars; GLM with corrected Tukey’s multiple comparisons post-hoc; p<0.0001) and the A8 (86%; 1029 synapses: 1202 total synapses; white bars; GLM with corrected Tukey’s multiple comparisons post-hoc; p<0.001). Thus, the relative frequency of CRF-positive synaptic terminals on TH-positive versus TH-negative profiles was similar in the PBP and A8 (within TH-positive comparisons: p=0.3741 n.s.; within TH-negative comparisons: p=0.0817 n.s., GLM with corrected Tukey’s multiple comparisons test).

**Figure 6.**
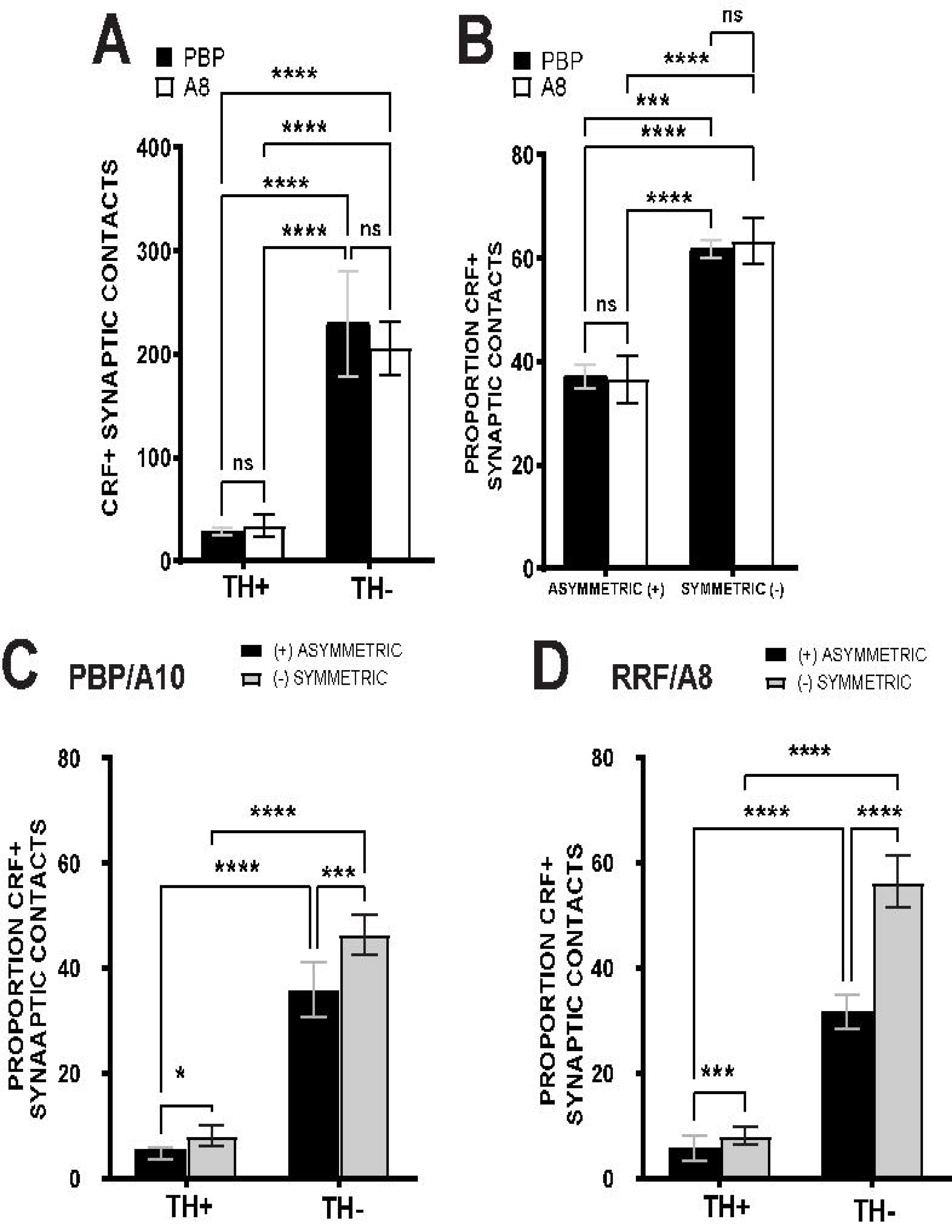
CRF positive axon contacts in PBP/A10 and RRF/A8. (**A**) CRF- positive axon synapses across TH- positive versus TH-negative profiles in the PBP and A8. Significantly more synapses were found on TH- negative profiles in both PBP and A8. Between region comparisons were not significantly different. (**B**) Relative proportion of CRF synapse type (asymmetric,(+) excitatory; symmetric,(-) inhibitory) in PBP and A8. There are significantly more symmetric than asymmetric synapses in both PBP and A8, with no significant differences between regions. (**C**) Proportion of CRF-positive synaptic contacts by synapse type (asymmetric (+) vs symmetric (-)) onto TH-positive and TH-negative profiles in the PBP/A10. Significantly more CRF-positive synapses (both asymmetric and symmetric) were found on TH-negative profiles. The proportion of asymmetric versus symmetric synapses was significantly increased on both TH-positive versus TH-negative profiles. (**D**) Proportion of CRF-positive synaptic contacts across TH-positive and TH- negative profiles, and the relative proportion of synapse type onto each group (asymmetric (+) vs symmetric (-)) in RRF/A8. As in the PBP, significantly more synapses (both asymmetric and symmetric) were found in TH-negative profiles. Within each category of post-synaptic partners, the relative proportion of symmetric synapses was significantly higher than asymmetric synapses. *= p≤ 0.05. **= p<0.005, ***= p<0.001, ****= p<0.0001.

### The proportion of CRF-positive terminals with inhibitory synapses is greater than those with excitatory synapses in both PBP and A8

We next quantified the proportion of either asymmetric (excitatory) or symmetric (inhibitory) synapses within each region (**Figure 6B**). In the PBP and A8, symmetric synapses comprised the majority of all synapses (62% in the PBP, black bars, p=0.0008, Tukey’s multiple comparisons test; 63% in the A8, A8, white bars, p=0.0001, Tukey’s multiple comparisons test). Within each synaptic profile group (asymmetric or symmetric), no differences were found in each anatomical region (PBP or A8) suggesting that a similar proportion of each synaptic profile were found (A8-asymmetric: PBP-asymmetric, p=0.9999; A8- symmetric: PBP-symmetric, p=0.9963).

### Characterizing type of CRF synapse onto TH-positive and TH-negative profile types in the PBP and A8

We next investigated the nature of CRF-immunoreactive synaptic contacts interactions by quantifying the proportion of CRF-positive axon synapses of both types (asymmetric [+] vs symmetric [-]) across both dendrite types (TH+ vs TH-) in each region. In the PBP (**Figure 6C**), there were considerably more synaptic contacts overall onto TH-negative profiles compared to TH-positive profiles. Because of the significant weighting of CRF synapses on TH-negative dendrites overall, the vast majority of both asymmetric and symmetric-type synapses were also found on these dendritic profiles. 89% of asymmetric contacts (black bars; 143 synapses: 162 total synapses) were found on TH-negative profiles (p<0.0001, GLM with corrected Tukey’s multiple comparisons tests), and 85% of symmetric synapses (gray bars; 185 synapses: 217 total synapses) were found on TH-negative profiles (p<0.0001, GLM with corrected Tukey’s multiple comparisons test). Within each of the cell groups (TH-positive and TH-negative) there were significantly more symmetric contracts when compared to asymmetric synapses (TH-positive, asymmetric [black bar] vs symmetric [gray bar], p=0.281; TH-negative, asymmetric [black bar] vs symmetric [gray bar], p=0.0001).

In the RRF/A8 (**Figure 6D**), we also found considerably more overall synaptic contacts onto TH-negative profiles compared to TH-positive profiles. For the asymmetric synapses (black bars), 85% were onto TH- negative profiles (158 total synapses: 186 total overall synapses; p<0.0001; Tukey’s multiple comparisons test). Among the symmetric synapses (gray bars), we found that 88% of those synapses were also onto TH-negative profiles (281 total synapses: 321 overall synapses; p<0.0001, Tukey’s multiple comparisons test). We next analyzed differences in the proportion of asymmetric versus symmetric synapses across post-synaptic profile types (TH-positive vs TH-negative). TH-negative profiles had significantly greater proportions of symmetric (inhibitory, gray bar) type synapses compared to excitatory type synapses (black bar, p<0.0001). Similarly, there were significantly more symmetric (gray bar) type synapse compared to asymmetric (black) type synapses in the TH-positive profiles in RRF/A8.

### Frequency of CRF-positive synaptic contacts are comparable across males and females

CRF is differentially regulated in males and females via several mechanisms (Bangasser and Valentino 2012; Bangasser et al. 2013). In females, stress induces enhanced CRF-mediated activation of the HPA axis and has enhanced effects on post-synaptic receptor dynamics (reviewed in, Bangasser and Valentino 2012). Therefore we compared CRF-positive synaptic contacts across these profile in males (blue) and females (red) (**Figure 7**). Similar to pooled cases (**Figure 6**), in the PBP, we noted significantly more synapses made on TH-negative profiles in males, with 93% of all synapses on TH-negative profiles (**Figure 7A**, solid blue bars,585 synapses: 631 total synapses, p<0.0001, corrected Tukey’s multiple comparisons tests). In females, 83% of all synapses were on TH-negative profiles in the PBP (**Figure 7A**, solid red bars, 332 synapses: 400 total synapses, p<0.0001, corrected Tukey’s multiple comparisons tests). There were thus no significant gender differences between the total number of CRF-positive contacts onto relatively low numbers of TH-positive profiles, but males had significantly more contacts onto TH-negative profiles compared to females (p<0.0001, corrected Tukey’s multiple comparisons tests).

**Figure 7.**
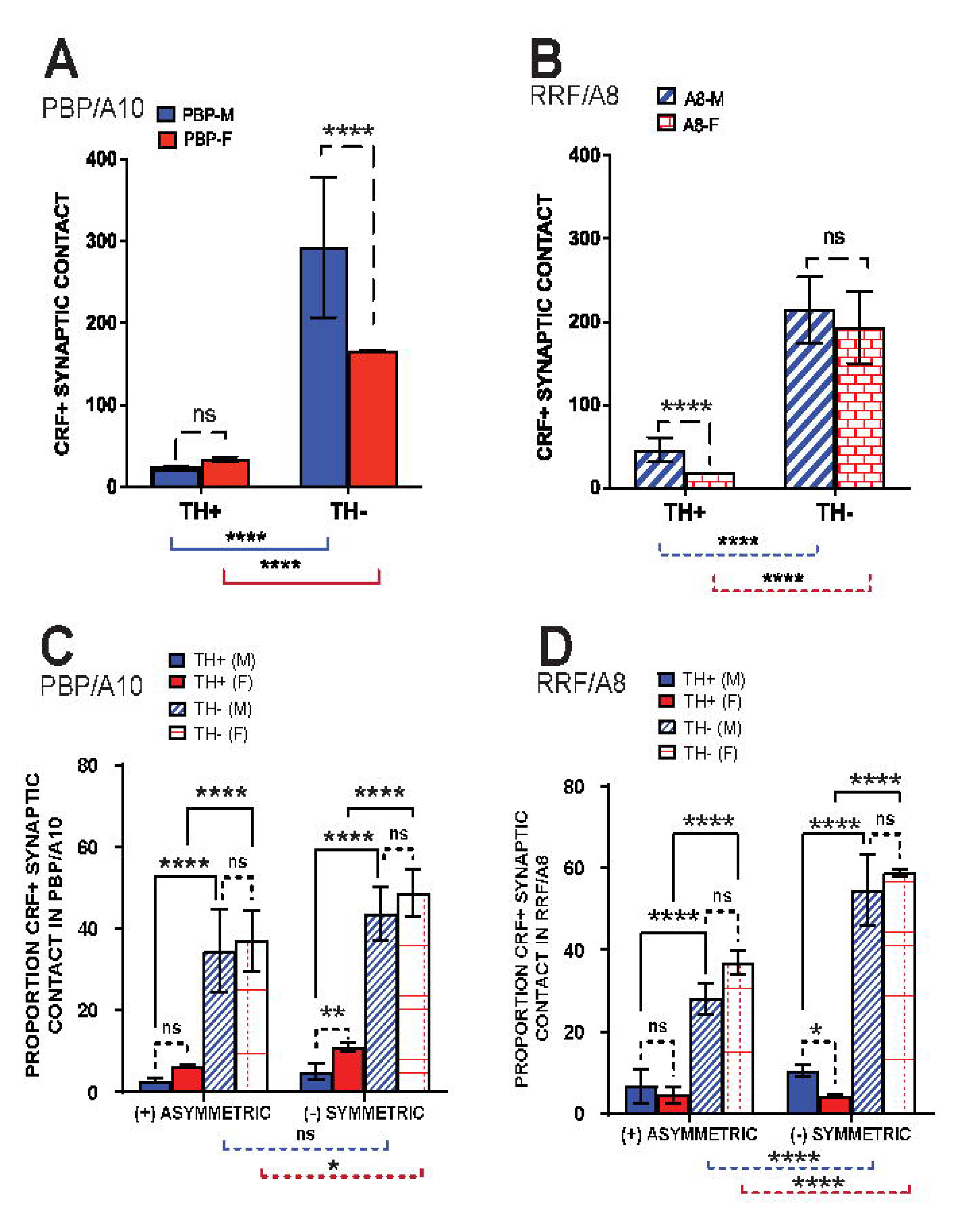
CRF-positive synaptic contacts in PBP/A10 and RRF/A8 in males vs females. (**A**, **B**) Comparison of all CRF-positive synapses onto TH-positive versus TH-negative post-synaptic profiles in the PBP (**A**) and A8 (**B**) in males and females. (**C-D**) Proportion of CRF-IR asymmetric and symmetric contacts onto TH- positive and TH-negative dendrites in the PBP (**C**) and A8 (**D**), considered in males (blue) and females (red). There were no differences in the proportion of asymmetric (+) versus symmetric (-) type synapses between males and females. Proportions of both types of CRF-positive synapses were significantly higher on the TH-negative (compared to TH-positive) profiles, with no differences between in males versus females. In both PBP (C) and A8 (D), there was a significantly higher proportion of total asymmetric-type synapses versus symmetric synapses overall. Significantly more symmetric synapses were found on TH- negative profiles in both sexes (male: blue comparison bar below graph, female: red comparison bar below graph). See text for details. *= p≤ 0.05. **= p<0.005, ***= p<0.001, ****= p<0.0001.

In RRF/A8, 86% of all male CRF-positive synapses were onto TH-negative profiles (**Figure 7B**, hatched blue bars, 462 synapses: 538 total synapses, p<0.0001), and that 91% of all female CRF-positive synapses onto TH-negative profiles (387 synapses: 423 total synapses, hatched red bars, p,0.0001). In contrast to the PBP, in the RRF/A8, males had significantly more CRF-labeled contacts onto TH-positive profiles compared to females (blue hatched bar vs red hatched bar, p<0.0001), while CRF-positive contacts onto TH-negative profiles were similar across the sexes. Thus, while general patterns of greater numbers of CRF immunoreactive synaptic contacts onto TH-negative versus TH-positive dendritic profiles held for both sexes, there were differences in the relative numbers of these synaptic contacts for each sex.

We next compared the proportion of synapse type (asymmetric and symmetric) across TH-positive and TH- negative profiles in males and females for each region. In the PBP (**Figure 7C**), for both sexes, there were more CRF-positive synapses onto TH-negative versus TH-profiles regardless of synapse type. In males, 92% of all asymmetric synapses were onto TH-negative profiles (69 synapses: 75 total asymmetric synapses, p<000.1), whereas, 89% of all symmetric synapses were onto TH-negative profiles (87 synapses: 97 total symmetric synapses, p<000.1) compared to TH-positive profiles. In females, 85% of all asymmetric synapses were onto TH-negative profiles (solid red vs hatched red, TH-negative asymmetric: p<0.0001; 74 synapses: 87 total asymmetric synapses) and 81% of all symmetric synapses were onto TH- negative profiles (solid red vs hatched red, TH-negative symmetric: p<0.0001; 98 synapses:120 total symmetric synapses). We did not find significant sex differences between sexes when comparing within synapse type for most comparisons (blue solid vs red solid; hatched blue vs hatched red, for all comparisons p>0.9999), except in one comparison (TH-positive symmetric synapses in males [solid blue bar] vs TH-positive symmetric synapses in females [solid red bar], p=0.0021). Given the disproportionate number of synapses onto TH-negative profiles (versus TH-positive profiles) in PBP, we further compared whether there were more asymmetric vs symmetric synapses within this subpopulation of profiles. While no significant differences were found in males (blue hatched bars, p=0.9146), the proportion of symmetric synapses in TH-negative profiles was significantly higher compared to asymmetric synapses in females (red hatched bars, p=0.0195).

Similarly, in RRF/A8 (**Figure 7D**), there was a general pattern of more CRF-positive synapses onto TH- negative versus TH-profiles regardless of synapse type regardless of sex. In males, we found 81% and 84% of all asymmetric (84 synapses: 104 total synapses) and symmetric synapses (164 synapses: 195 total synapses), respectively, onto TH-negative profiles compared to TH-positive profiles. In across-cell type comparisons, males showed a significant difference in the proportion of asymmetric synapses on TH- positive vs TH-negative profiles (solid blue bar vs hatched blue bar, p<0.001). Similarly, significantly more symmetric synapses were made onto TH-negative profiles compared to TH-positive profiles in males (blue bar vs hatched blue, p<0.001). Females showed a similar pattern with 89% of all asymmetric synapses (74 synapses: 83 total synapses) and 93% of all symmetric synapses (118 synapses: 126 total synapses) on TH-negative profiles. In across-cell type comparisons, significantly more CRF-positive contacts of both types were found on TH-negative profiles compared to TH-positive profiles (TH+ versus TH-, female asymmetric synapses, solid red bar vs hatched red bar, p<0.0001; female symmetric synapses, solid red bar vs hatched red bar, p<0.0001; Tukey’s multiple comparisons test). We did not see significant differences in male vs female proportions for most comparisons in RRF/A8, with the exception that the proportion of symmetric synapses onto TH-positive profiles in males was significantly higher in males (Male vs Female, symmetric synapses comparisons within TH-positive profiles, p=0.0174). To further assess the disproportionate number of synapses in TH-negative profiles in A8, we further compared whether there were more asymmetric vs symmetric synapses within this subpopulation. We found significantly more symmetric synapses across both sexes in TH-negative profiles (males, blue comparison bar below graph, p<0.0001; females, red comparison bar below graph, p<0.0001).

Overall, we show that both DA subregions (PBP and A8) had the highest proportion of CRF-positive synapses onto TH-negative profiles with the proportion of CRF-positive asymmetric vs symmetric synapses within each of these profiles significantly higher in the asymmetric synapse population. Our findings are consistent across both pooled (**Figure 6**) and sex comparison data (**Figure 7**) and suggest a unique CRF- mediated modulation of TH-negative (presumed GABAergic) profiles in the PBP/A10 and RRF/A8 in the macaque midbrain.

### Cortisol and sex hormones are stable and comparable over 6 months in young males and females

How CRF expression and stress impact the DA system may be developmentally regulated (Rincon-Cortes and Grace 2017; Izzo et al. 2005; Coco et al. 1992). Adolescent monkeys are transitioning into sexual maturation at three to five years of age, with changes in physical parameters (vaginal epithelial changes, testicular volume) as well as hormonal shifts (Plant 2015). To determine if pubertal status or stress effects was a likely factor in our cohort, we performed gonadal hormone and cortisol assays from serum specimens on seven animals (three of which are included in this study) at several timepoints (**Table 3**). In serum preparations, analyte concentrations were comparable between males and females with expected differences between gonadal hormones. Stress hormones in our pair-housed animals were not significantly different across animals (**Figure 8**). These findings suggest that, in our pair-housed animal environment, gonadal hormone and stress hormones are stable and comparable in same-aged animals over a 6-month time period.

**Table 3.**
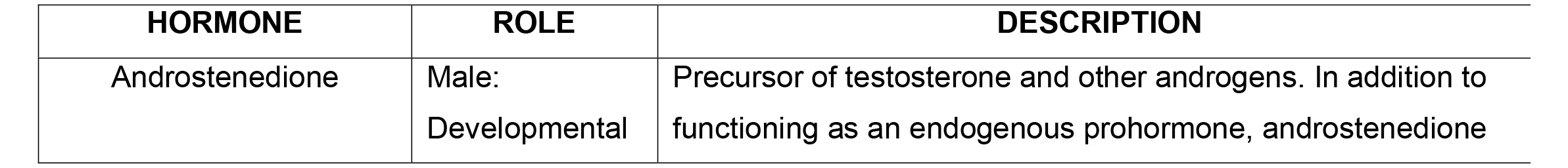
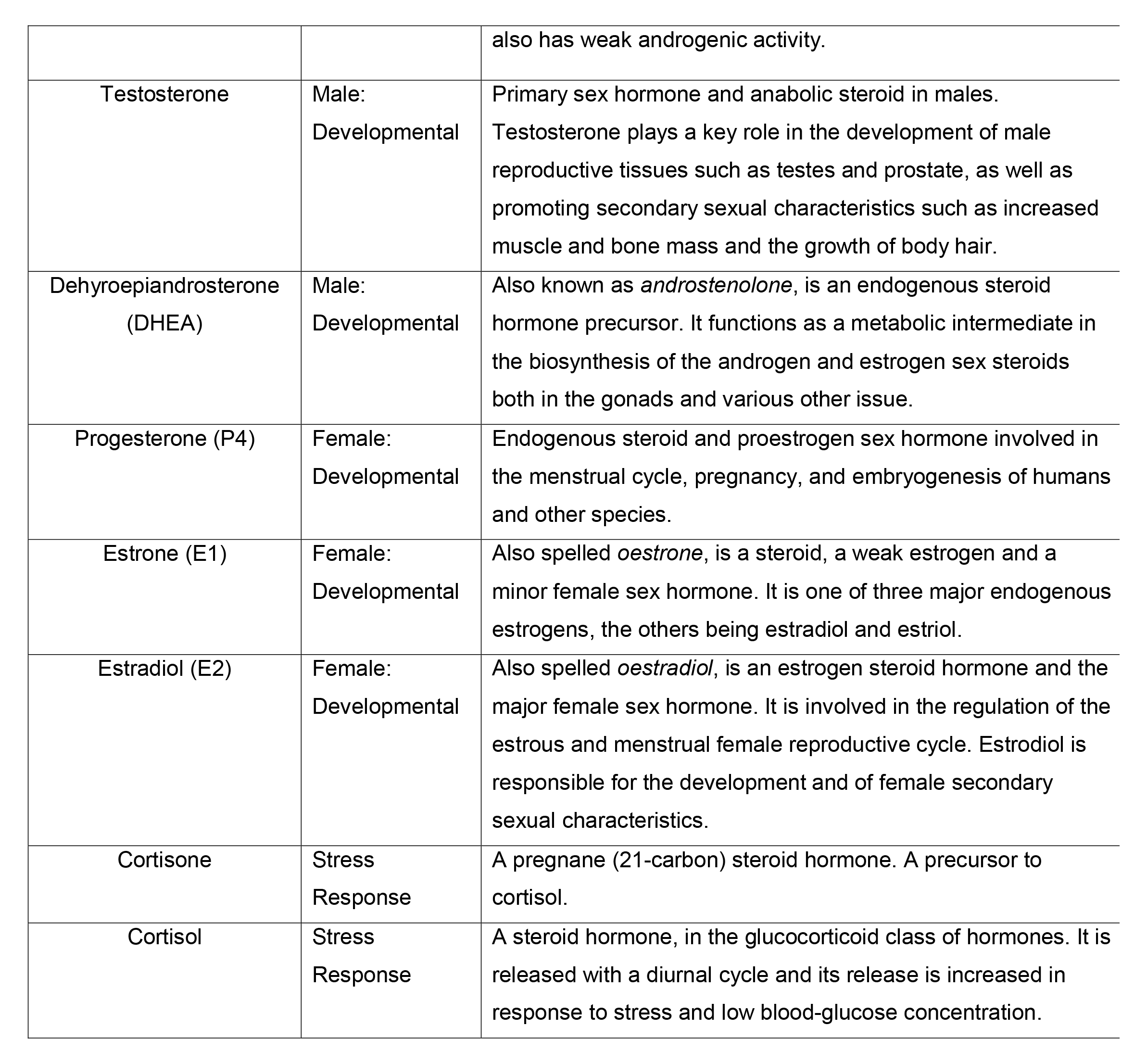

| **HORMONE** | **ROLE** | **DESCRIPTION** |
| --- | --- | --- |
| Androstenedione | Male:<br>Developmental | Precursor of testosterone and other androgens. In addition to functioning as an endogenous prohormone, androstenedione |
| Pubertal and stress hormone descriptions |  |  |
|  |  | also has weak androgenic activity. |
| Testosterone | Male:<br>Developmental | Primary sex hormone and anabolic steroid in males. Testosterone plays a key role in the development of male reproductive tissues such as testes and prostate, as well as promoting secondary sexual characteristics such as increased muscle and bone mass and the growth of body hair. |
| Dehydroepiandrosterone (DHEA) | Male:<br>Developmental | Also known as *androstenolone*, is an endogenous steroid hormone precursor. It functions as a metabolic intermediate in the biosynthesis of the androgen and estrogen sex steroids both in the gonads and various other issue. |
| Progesterone (P4) | Female:<br>Developmental | Endogenous steroid and proestrogen sex hormone involved in the menstrual cycle, pregnancy, and embryogenesis of humans and other species. |
| Estrone (E1) | Female:<br>Developmental | Also spelled *oestrone*, is a steroid, a weak estrogen and a minor female sex hormone. It is one of three major endogenous estrogens, the others being estradiol and estriol. |
| Estradiol (E2) | Female:<br>Developmental | Also spelled *oestradiol*, is an estrogen steroid hormone and the major female sex hormone. It is involved in the regulation of the estrous and menstrual female reproductive cycle. Estrodiol is responsible for the development and of female secondary sexual characteristics. |
| Cortisone | Stress<br>Response | A pregnane (21-carbon) steroid hormone. A precursor to cortisol. |
| Cortisol | Stress<br>Response | A steroid hormone, in the glucocorticoid class of hormones. It is released with a diurnal cycle and its release is increased in response to stress and low blood-glucose concentration. |

## DISCUSSION

In this study, we investigated the relationship between CRF and its terminal synapses on TH-positive versus TH-negative profiles in the PBP and A8 regions. These DA subregions were investigated since they are evolutionarily expanded in nonhuman primates (Halliday and Tork 1986; Fu et al. 2016; McRitchie et al. 1996; Francois et al. 1999), and are sites of intense CRF-immunoreactive fiber termination. Importantly, the PBP and A8 neurons have wide-ranging efferents which are different from those of the midline VTA (Fudge et al. 2017; Haber et al. 2000; Williams and Goldman-Rakic 1998; Francois et al. 1999)(see below). We found that (1) CRF-positive axons terminated mainly onto TH-negative profiles in the PBP and A8 DA subregions; (2) CRF-positive axon terminals formed both asymmetric and symmetric synapses, with a higher frequency of symmetric (inhibitory-type) synapses in both PBP and A8; (3) In both the PBP and A8, TH-negative dendritic profiles and TH-positive profiles each received significantly more inhibitory than excitatory-type CRF synapses, and (4) same age male and female subjects had similar findings on these measures, despite shifts in the relative numbers of CRF-positive synapses onto TH- positive and TH-negative across regions.

**Figure 8.**
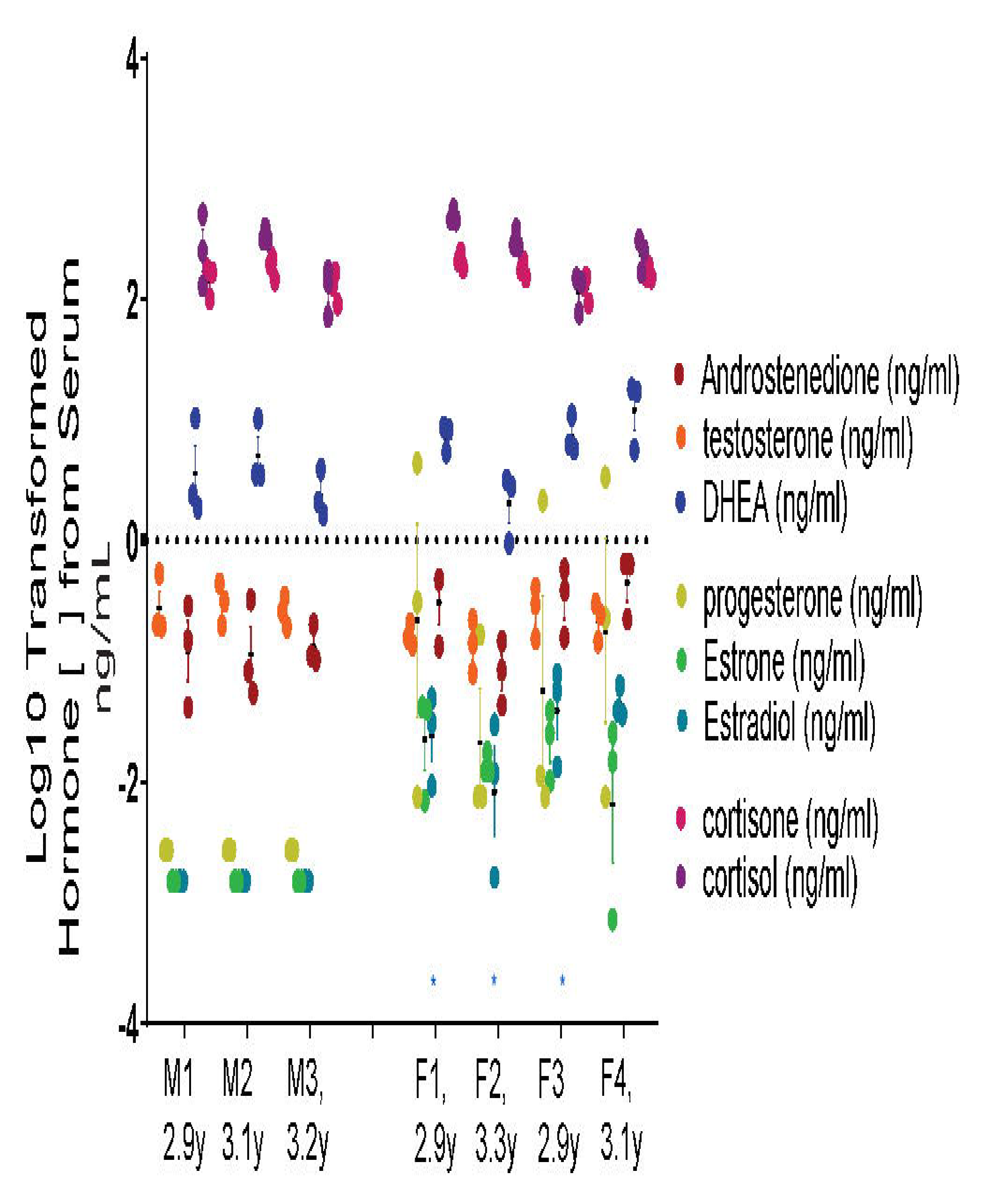
Pubertal and stress hormone assays in adolescent male and female macaques. Gender-specific developmental hormones and stress hormone concentrations from serum. Male hormones: androstenedione, testosterone, and DHEA (brown, orange, blue); Female hormones progesterone, estrone and estradiol (yellow, green, blue). Stress hormones: cortisone and cortisol (pink, purple). Asterisks show animals used in the present study. Descriptions of hormones can be found in **Table 3**.

Together, our results suggest that CRF is mainly expressed in GABAergic terminals, that largely contact TH-negative profiles in both PBP and A8. While we did not determine the phenotype of TH-negative profiles, we hypothesize that they are mostly GABAergic. Glutamatergic neurons also exist in the ventral midbrain, but are not a majority subpopulation (Margolis et al. 2012; Yamaguchi et al. 2013; Root et al. 2016; Nair-Roberts et al. 2008). They are highest in the rostrolinear nucleus of the midline VTA group, but drop off substantially in other subregions of the ventral midbrain, particularly in higher species. We thus hypothesize that the net effect of CRF/GABA synapses under basal conditions in the PBP and A8 is inhibition of post-synaptic GABAergic interneurons, which can release DA from inhibitory control in specific output paths (**Figure 9**).

**Figure 9.**
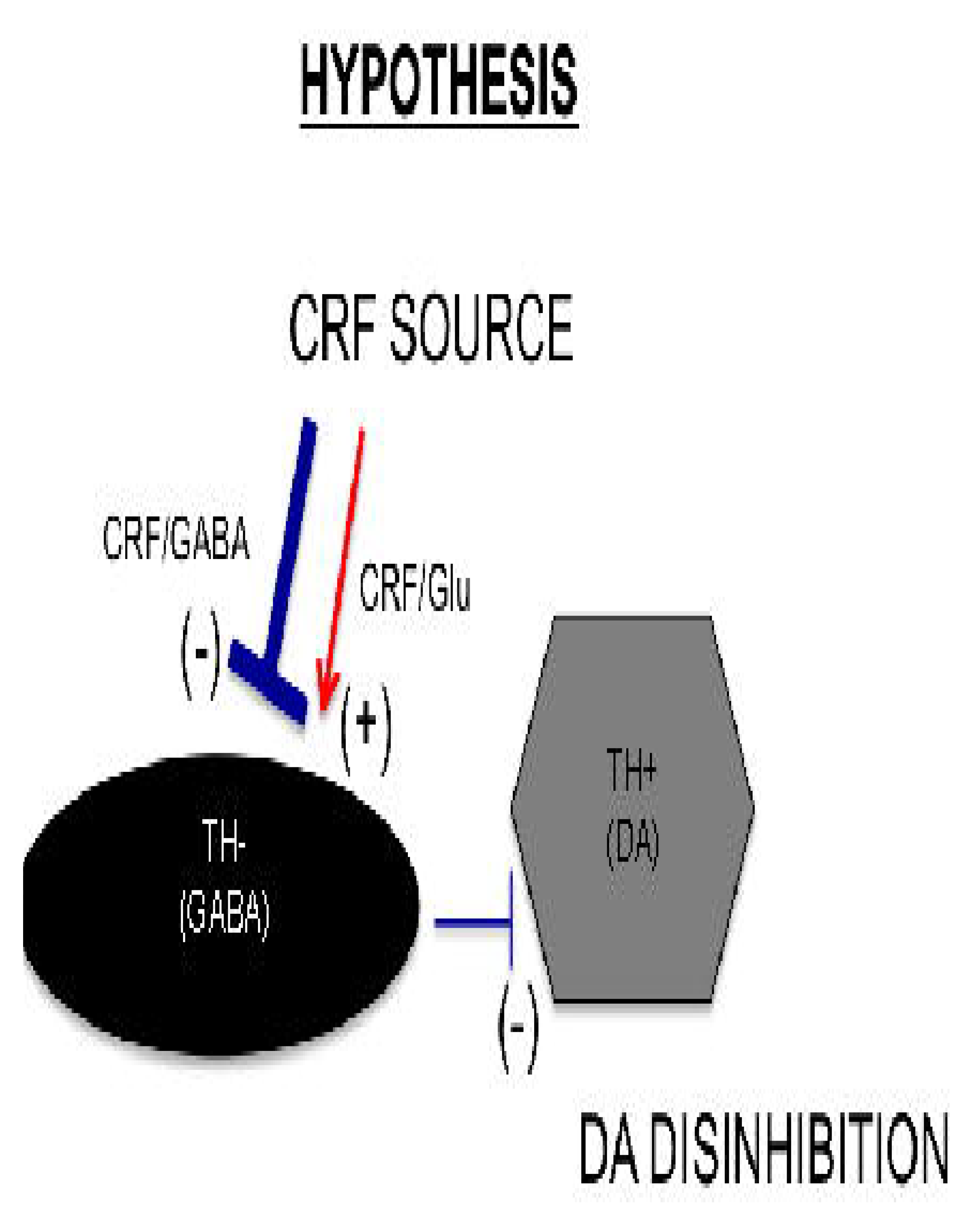
Indirect CRF modulation of DA cells. CRF-positive synaptic terminals predominantly make symmetric (inhibitory) synapses onto TH-negative profiles in both PBP and A8. We hypothesize that under stress, CRF in inhibitory synapses may enhance GABA effects on TH-negative profiles (presumptive GABA interneurons), resulting in a disinhibition of DA neurons.

### Comparison with previous work: CRF modulation of DA subcircuits

Surprisingly, only one study has examined CRF expression in excitatory and inhibitory synapses in the VTA using EM (Tagliaferro and Morales 2008). In that rodent study, which broadly examined all VTA subnuclei (including the PBP), the majority (58%) of CRF-IR axon terminals targeted non-DA cells, and were predominantly symmetric (inhibitory) and thus, GABAergic in nature. We found, similarly, that the majority of CRF-positive synapses were onto TH-negative profiles in the monkey PBP and A8 (**Figure 6**). In contrast to the Tagliaferro study, however, we did not find a preponderance of excitatory-type CRF profiles on TH-positive profiles in either the PBP or A8. In fact, the overall number of CRF-containing terminals onto TH-positive profiles were relatively small in both PBP and A8.

### Sources of CRF in the ventral midbrain

CRF fibers originate from multiple potential afferent sources to innervate the DA system (reviewed in, Kelly and Fudge 2018). Although the BSTL, CeN, and entire extended amygdala have many CRF-positive neurons and are important for stress-mediated behaviors, there are many potential sources of CRF- positive afferent inputs to the ventral midbrain (Kelly and Fudge 2018). Potential structures that project to the ventral midbrain and contain CRF-positive cells include: ventral striatum, midline thalamic nuclei, the periaqueductal gray (rodents only), the peripeduncular nucleus, the pedunculopontine tegmental nucleus, the parabrachial nucleus, the lateral dorsal tegmental nucleus, dorsal mesencephalic nucleus, the median raphe, and non-noradrenergic cells of the locus coeruleus. A detailed mapping study to determine whether CRF-containing neurons in these areas actually contribute CRF terminals to specific DA subregions is of interest.

Moreover, CRF innervation of DA subregions outside the midline VTA (A10) has generally been ignored. However Dabrowska et al. (2016) used genetic approaches in mice and rats to show that CRF- synthesizing cells in the BSTL send relatively more intense fiber labeling over the substantia nigra pars compacta compared to the VTA. In the nonhuman primate, using anterograde tracing methods, we similarly find that the BSTL and CeN send a relatively dense input *outside* the midline VTA subregion, i.e., to the PBP and A8. Validating this finding with retrograde tracer injections in the PBP and A8, we showed that a majority of tracer-labeled cells in the BSTL and CeN co-contained CRF protein (Fudge et al. 2017). Interestingly, our recent examination of CRF neuron phenotypes throughout the monkey BSTL, sublenticular extended amygdala, and CeN revealed that CRF mRNA is mainly expressed against a background of vesicular GABA transporter (VGAT), but that there are a number of ‘multiplexed’ CRF- labeled neurons that co-contain both VGAT and vesicular glutamatergic transporter 2 (VGluT2), depending on subregion (Fudge et al. 2022). The CRF-labeled neurons in the BSTL to CeN continuum (known as the ‘extended amygdala’) thus have a mixture of both ‘pure’ GABA and ‘multiplexed’ (GABA/glutamate) phenotypes. While only one source of CRF to the PBP and A8, this co-transmitter profile is generally consistent with overall balance of inhibitory versus excitatory CRF synapses found in this study. It will be important to conduct cell-type specific tracer studies, to precisely ascertain whether the CRF-extended amygdala to PBP/A8 path arises mainly from GABAergic (inhibitory) neurons, and the relative contribution of ‘multiplexed’ neurons.

### Balance of DA and GABA neurons across PBP and A8: implications for CRF synaptic data

An important consideration in assessing quantitative data on CRF terminals pertains to the difference in ratios of DA:GABA post-synaptic partners in each region. GABAergic neurons comprise the largest non-dopaminergic cell population in the ventral midbrain (Nair-Roberts et al. 2008; Margolis et al. 2012), and contribute to potential modulation of local (and possible long-range) DA circuits (Omelchenko and Sesack 2009; Carr and Sesack 2000). In monkey, the ratio of DA to non-DA cells varies across the A10, A9 and A8 groups, and differs from the ratios in rodents (Poirier et al. 1983; German and Manaye 1993; Nair-Roberts et al. 2008; Swanson 1982). We recently quantified the distribution of DA neurons and their GABAergic counterparts throughout A10 (midline VTA and PBP separately), A9 (SNc) and A8 (RRF) subregions in the males of the present cohort using unbiased stereology (Kelly et al. 2022) and found significant regional differences in the balance of DA and GABAergic neurons. While there was an overall 3-fold greater number of DA neurons compared to GABAergic neurons, sub-regional ratios varied widely. The PBP has almost 5 times more DA neurons than GABAergic neurons; in contrast, in the A8 region, the ratio is close to 1:1. Here, we find that CRF fibers predominantly targeted TH-negative (presumptive GABAergic neurons) in both PBP and A8, forming symmetric (inhibitory) synapses. Under our arbitrary sampling approach, there was an equal probability of finding a CRF-containing synapse on a TH-positive versus TH-negative profile. However, in the PBP, where DA neurons outnumbered GABA neurons by 5:1, there presumably was a smaller probability of finding CRF-TH-negative synapses. Therefore, we suspect that in the PBP, we may be underestimating TH-negative/CRF terminal interactions, where in the A8 region, where GABA and DA neuron ratios are 1:1, estimates of CRF synapses are likely more accurate.

### Non-synaptic CRF

CRF release at the synapse also has 'parasynaptic' effects (Hokfelt et al. 2003). In other words, each time CRF is synaptically released, it is capable of diffusing long distances, and has complementary effects on other cells in the region. Therefore, there are important, additional modulatory effects of CRF on post-synaptic cells and other cells in the region, some of which are mediated at CRF1 and CRF2 receptors. Diffusion effects on distant neurons are best understood electrophysiologically, but the interpretation of parasynaptic effects will certainly be aided by understanding where CRF-positive synapses are located in the midbrain, the co-transmitter profiles associated with them, and the identity of their post-synaptic partners.

### DA and stress: beyond the classic VTA

During stress exposure in rodents, CRF is released into the VTA in an activity dependent manner (Wang et al. 2005). CRF’s postsynaptic effects on DA neurons can be inhibitory (Beckstead et al. 2009) or excitatory (Ungless et al. 2003; Wanat et al. 2008), and is pathway specific (Wanat et al. 2013). While CRF has little effect on cellular activity under non-stress conditions, chemogenetic manipulation of CRF-expressing GABA neurons in the CeN shows that CRF is critical for enhancing fear acquisition (Pomrenze et al. 2019). These findings are consistent with the idea that CRF serves as a behavioral amplifier under stressful conditions by enhancing the action of the primary transmitter on post-synaptic neurons. Our results with respect to the DA system show that CRF terminals from all sources in the PBP and A8 mainly form inhibitory-type synapses onto TH-negative (presumptive GABAergic) neurons, and may act to indirectly enhance DA signaling in stressful situations.

DA is a key player in modulating diverse functions through specific input/output pathways. While initially viewed as a 'syncytium' (or mass) of cells, the midbrain DA system is now recognized to be highly heterogeneous with respect to both circuits and function (Davis et al. 1991; Brown et al. 1979). Elegant models in rodents increasingly support this view, showing that the adjacent regions of the VTA are differentially involved in circuits and selective behaviors (de Jong et al. 2019; Steinberg et al. 2020; Menegas et al. 2018). The ways that different forms of stress affect specific DA subcircuits is a relatively new frontier (Kong and Zweifel 2021; Verharen et al. 2020; Douma and de Kloet 2020).

Electrophysiologic studies in monkeys would suggest that PBP and A8 subregions of the ventral midbrain—in contrast to the midline VTA-- largely signal the ‘salience’ of stimuli, regardless of reward value (Bromberg-Martin et al. 2010). These more laterally placed DA neurons, which coincide anatomically with the PBP and A8, fire to ‘non-reward’ stimuli that are nonetheless salient, and are important for orienting and heightened learning during unpredictable cues (Lee et al. 2006; Menegas et al. 2017). Here we find that these laterally located DA subregions are enriched in CRF-containing fibers, which may be important in shaping behavior responses to unexpected, salient information. Consistent these functions, DA neurons in the PBP and A8 in monkey project to the central (‘associative’) striatum (Fudge et al. 2017; Haber et al. 2000) and also the prefrontal cortex(Williams and Goldman-Rakic 1998). CRF-contacts on the non-dopaminergic neurons of the PBP (‘lateral’ VTA) and A8, suggest a novel mode for tuning DA responses in output paths associated with cognitive functions in higher species, a mechanism that we hypothesize is important for responses to unpredictable cues and uncertainty in health and disease states (Horga and Abi-Dargham 2019; Balsters et al. 2020; Hupalo et al. 2019).

## STATEMENTS & DECLARATIONS

### Author Contributions

All authors contributed to the study design. Material preparation, data collection and analysis were performed by Emily A. Kelly and Tanzy Love. The first draft and all additional drafts of the manuscript were written by Emily A. Kelly and Julie L. Fudge. All authors read and approved the final manuscript.

### Data Availability

Data and materials supporting the results or analyses presented in this study are available upon reasonable request from the corresponding author.

### Ethics approval

All experiments were conducted in accordance with National Institute of Health guidelines (NIH Publications No. 80-23). Experimental design and technique were aimed at minimizing animal use and suffering and were reviewed by the University of Rochester Committee on Animal Research.

## ACKNOWLEDGEMENTS

We’d like to thank the URMC Electron Microscope Research Core for their expert guidance and assistance, Nanette Alcock (University of Rochester SMD) for histological preparations, Amita Kapoor (The Wisconsin National Primate Research Center, University of Wisconsin-Madison) for hormone assay processing (P51OD011106) and funding by the National Institutes for Mental Health (RO1MH115016, J.L.F.)

## REFERENCES

Arias-Carrion O, Stamelou M, Murillo-Rodriguez E, Menendez-Gonzalez M, Poppel E (2010) Dopaminergic reward system: a short integrative review. Int Arch Med 3:24. doi:10.1186/1755-7682-3-24

Arsenault MY, Parent A, Seguela P, Descarries L (1988) Distribution and morphological characteristics of dopamine-immunoreactive neurons in the midbrain of the squirrel monkey (Saimiri sciureus). J Comp Neurol 267:489–506

Balsters JH, Zerbi V, Sallet J, Wenderoth N, Mars RB (2020) Primate homologs of mouse cortico-striatal circuits. eLife 9. doi:10.7554/eLife.53680

Bangasser DA, Reyes BA, Piel D, Garachh V, Zhang XY, Plona ZM, Van Bockstaele EJ, Beck SG, Valentino RJ (2013) Increased vulnerability of the brain norepinephrine system of females to corticotropin-releasing factor overexpression. Molecular psychiatry 18 (2):166–173. doi:10.1038/mp.2012.24

Bangasser DA, Valentino RJ (2012) Sex differences in molecular and cellular substrates of stress. Cellular and molecular neurobiology 32 (5):709–723. doi:10.1007/s10571-012-9824-4

Beckstead MJ, Gantz SC, Ford CP, Stenzel-Poore MP, Phillips PE, Mark GP, Williams JT (2009) CRF enhancement of GIRK channel-mediated transmission in dopamine neurons. Neuropsychopharmacology: official publication of the American College of Neuropsychopharmacology 34 (8):1926-1935. doi:10.1038/npp.2009.25

Beier KT, Steinberg EE, DeLoach KE, Xie S, Miyamichi K, Schwarz L, Gao XJ, Kremer EJ, Malenka RC, Luo L (2015) Circuit Architecture of VTA Dopamine Neurons Revealed by Systematic Input-Output Mapping. Cell 162 (3):622–634. doi:10.1016/j.cell.2015.07.015

Bjorklund A, Dunnett SB (2007) Dopamine neuron systems in the brain: an update. Trends in neurosciences 30 (5):194–202. doi:S0166-2236(07)00067-7 [pii] 10.1016/j.tins.2007.03.006

Bouvier D, Corera AT, Tremblay ME, Riad M, Chagnon M, Murai KK, Pasquale EB, Fon EA, Doucet G (2008) Pre-synaptic and post-synaptic localization of EphA4 and EphB2 in adult mouse forebrain. J Neurochem 106 (2):682–695. doi:10.1111/j.1471-4159.2008.05416.x

Bromberg-Martin ES, Matsumoto M, Hikosaka O (2010) Dopamine in motivational control: rewarding, aversive, and alerting. Neuron 68 (5):815–834. doi:10.1016/j.neuron.2010.11.022

Brown RM, Crane AM, Goldman PS (1979) Regional distribution of monoamines in the cerebral cortex and subcortical structures of the rhesus monkey: concentrations and in vivo synthesis rates. Brain research 168 (1):133–150

Carr DB, Sesack SR (2000) GABA-containing neurons in the rat ventral tegmental area project to the prefrontal cortex. Synapse 38 (2):114–123. doi:10.1002/1098-2396(200011)38:2<114::AID-SYN2>3.0.CO;2-R

Cho YT, Fudge JL (2010) Heterogeneous dopamine populations project to specific subregions of the primate amygdala. Neuroscience 165 ():1501-1518. doi:S0306-4522(09)01800-4 [pii] 10.1016/j.neuroscience.2009.11.004

Coco ML, Kuhn CM, Ely TD, Kilts CD (1992) Selective activation of mesoamygdaloid dopamine neurons by conditioned stress: attenuation by diazepam. Brain research 590 (1-2):39–47

Cote P-Y, Sadikot AF, Parent A (1991) Complementary distribution of calbindin D-28k and parvalbumin in the basal forebrain and midbrain of the squirrel monkey. European Journal of Neuroscience 3:1316–1329

Craig AS (1974) Sodium-Borohydride as an Aldehyde Blocking Reagent for Electron-Microscope Histochemistry. Histochemistry 42 (2):141–144. doi:Doi 10.1007/Bf00533265

Dabrowska J, Martinon D, Moaddab M, Rainnie DG (2016) Targeting corticotropin-releasing factor (CRF) projections from the oval nucleus of the BNST using cell-type specific neuronal tracing studies in mouse and rat brain. Journal of neuroendocrinology. doi:10.1111/jne.12442

Davis KL, Kahn RS, Ko G, Davidson M (1991) Dopamine in schizophrenia: A review and reconceptualization. Am J Psychiatry 148:11–10

de Jong JW, Afjei SA, Pollak Dorocic I, Peck JR, Liu C, Kim CK, Tian L, Deisseroth K, Lammel S (2019) A Neural Circuit Mechanism for Encoding Aversive Stimuli in the Mesolimbic Dopamine System. Neuron 101 (1):133-151 e137. doi:10.1016/j.neuron.2018.11.005

Dosemeci A, Weinberg RJ, Reese TS, Tao-Cheng JH (2016) The Postsynaptic Density: There Is More than Meets the Eye. Front Synaptic Neurosci 8:23. doi:10.3389/fnsyn.2016.00023

Douma EH, de Kloet ER (2020) Stress-induced plasticity and functioning of ventral tegmental dopamine neurons. Neuroscience and biobehavioral reviews 108:48–77. doi:10.1016/j.neubiorev.2019.10.015

Farassat N, Costa KM, Stojanovic S, Albert S, Kovacheva L, Shin J, Egger R, Somayaji M, Duvarci S, Schneider G, Roeper J (2019) In vivo functional diversity of midbrain dopamine neurons within identified axonal projections. eLife 8. doi:10.7554/eLife.48408

Francois C, Yelnik J, Tande D, Agid Y, Hirsch EC (1999) Dopaminergic cell group A8 in the monkey: anatomical organization and projections to the striatum. Journal of Comparative Neurology 414 (3):334–347

Fu Y, Paxinos G, Watson C, Halliday GM (2016) The substantia nigra and ventral tegmental dopaminergic neurons from development to degeneration. Journal of chemical neuroanatomy 76 (Pt B):98–107. doi:10.1016/j.jchemneu.2016.02.001

Fudge JL, Kelly EA, Hackett TA (2022) Corticotropin releasing factor (CRF) co-expression in GABAergic, glutamatergic and GABA/glutamatergic subpopulations in the central extended amygdala and ventral pallidum of young male primates. The Journal of neuroscience: the official journal of the Society for Neuroscience. doi:10.1523/JNEUROSCI.1453-22.2022

Fudge JL, Kelly EA, Pal R, Bedont JL, Park L, Ho B (2017) Beyond the Classic VTA: Extended Amygdala Projections to DA-Striatal Paths in the Primate. Neuropsychopharmacology: official publication of the American College of Neuropsychopharmacology 42 (8):1563–1576. doi:10.1038/npp.2017.38

Gallagher JP, Orozco-Cabal LF, Liu J, Shinnick-Gallagher P (2008) Synaptic physiology of central CRH system. European journal of pharmacology 583 (2-3):215–225. doi:10.1016/j.ejphar.2007.11.075

Garcia JP, Keen KL, Kenealy BP, Seminara SB, Terasawa E (2018) Role of Kisspeptin and Neurokinin B Signaling in Male Rhesus Monkey Puberty. Endocrinology 159 (8):3048–3060. doi:10.1210/en.2018-00443

Gaspar P, Berger B, Gay M, Hamon M, Cesselin F, Vigny A, Javoy-Agid F, Agid Y (1983) Tyrosine hydroxylase and methionine-enkephalin in the human mesencephalon. J Neurol Sci 58:247–267

Gaspar P, Heizmann CW, Kaas JH (1993) Calbindin D-28K in the dopaminergic mesocortical projection of a monkey (Aotus trivirgatus). Brain Res 603:166–172

German DC, Manaye KF (1993) Midbrain dopaminergic neurons (nuclei A8,A9, and A10): three-dimensional reconstruction in the rat. J Comp Neurol 331:297-309

Grieder TE, Herman MA, Contet C, Tan LA, Vargas-Perez H, Cohen A, Chwalek M, Maal-Bared G, Freiling J, Schlosburg JE, Clarke L, Crawford E, Koebel P, Repunte-Canonigo V, Sanna PP, Tapper AR, Roberto M, Kieffer BL, Sawchenko PE, Koob GF, van der Kooy D, George O (2014) VTA CRF neurons mediate the aversive effects of nicotine withdrawal and promote intake escalation. Nature neuroscience 17 (12):1751–1758. doi:10.1038/nn.3872

Haber SN, Fudge JL (1997) The primate substantia nigra and VTA: Integrative circuitry and function. Crit Rev Neurobiol 11(4):323–342

Haber SN, Fudge JL, McFarland N (2000) Striatonigrostriatal pathways in primates form an ascending spiral from the shell to the dorsolateral striatum. J Neurosci 20:2369–2382

Halliday GM, Tork I (1986) Comparative anatomy of the ventromedial mesencephalic tegmentum in the rat, cat, monkey and human. J Comp Neurol 252:423–445

Henckens MJ, Deussing JM, Chen A (2016) Region-specific roles of the corticotropin-releasing factor-urocortin system in stress. Nature reviews Neuroscience 17 (10):636–651. doi:10.1038/nrn.2016.94

Hokfelt T, Bartfai T, Bloom F (2003) Neuropeptides: opportunities for drug discovery. Lancet Neurol 2 (8):463–472. doi:10.1016/s1474-4422(03)00482-4

Horga G, Abi-Dargham A (2019) An integrative framework for perceptual disturbances in psychosis. Nature reviews Neuroscience 20 (12):763–778. doi:10.1038/s41583-019-0234-1

Hupalo S, Bryce CA, Bangasser DA, Berridge CW, Valentino RJ, Floresco SB (2019) Corticotropin-Releasing Factor (CRF) circuit modulation of cognition and motivation. Neuroscience and biobehavioral reviews 103:50–59. doi:10.1016/j.neubiorev.2019.06.010

Izzo E, Sanna PP, Koob GF (2005) Impairment of dopaminergic system function after chronic treatment with corticotropin-releasing factor. Pharmacology, biochemistry, and behavior 81 (4):701–708. doi:10.1016/j.pbb.2005.04.017

Kalivas PW, Duffy P, Latimer LG (1987) Neurochemical and behavioral effects of corticotropin-releasing factor in the ventral tegmental area of the rat. The Journal of pharmacology and experimental therapeutics 242 (3):757–763

Kelly EA, Contraras JM, Duan A, Vassell R, Fudge JL (2022) Unbiased stereological estimates of dopaminergic and GABAergic neurons in the A10, A9 and A8 subpopulations in the young male Macaque. Neuroscience 496 (496):152-164. doi:doi.org/10.1016/j.neuroscience.2022.06.018

Kelly EA, Fudge JL (2018) The neuroanatomic complexity of the CRF and DA systems and their interface: What we still don’t know. Neuroscience and biobehavioral reviews. doi:10.1016/j.neubiorev.2018.04.014

Kelly EA, Tremblay ME, Gahmberg CG, Tian L, Majewska AK (2014) Subcellular localization of intercellular adhesion molecule-5 (telencephalin) in the visual cortex is not developmentally regulated in the absence of matrix metalloproteinase-9. The Journal of comparative neurology 522 (3):676–688. doi:10.1002/cne.23440

Kenealy BP, Keen KL, Kapoor A, Terasawa E (2016) Neuroestradiol in the Stalk Median Eminence of Female Rhesus Macaques Decreases in Association With Puberty Onset. Endocrinology 157 (1):70–76. doi:10.1210/en.2015-1770

Kong MS, Zweifel LS (2021) Central amygdala circuits in valence and salience processing. Behavioural brain research 410:113355. doi:10.1016/j.bbr.2021.113355

Lammel S, Ion DI, Roeper J, Malenka RC (2011) Projection-specific modulation of dopamine neuron synapses by aversive and rewarding stimuli. Neuron 70 (5):855–862. doi:10.1016/j.neuron.2011.03.025

Lammel S, Lim BK, Malenka RC (2014) Reward and aversion in a heterogeneous midbrain dopamine system. Neuropharmacology 76 Pt B:351-359. doi:10.1016/j.neuropharm.2013.03.019

Lavoie B, Parent A (1991) Dopaminergic neurons expressing calbindin in normal and parkinsonian monkeys. Neuroreport 2, No. 10:601–604

Lee HJ, Youn JM, O MJ, Gallagher M, Holland PC (2006) Role of substantia nigra-amygdala connections in surprise-induced enhancement of attention. The Journal of neuroscience: the official journal of the Society for Neuroscience 26 (22):6077–6081

Lewis DA, Sesack SR (1997) Dopamine systems in the primate brain. In: F.E. Bloom ABaTH (ed) Handbook of Chemical Neuroanatomy, vol. 13: The Primate Nervous System, Part 1, vol 13. Elsevier Science B.V., pp 263–375

Maley BE (1990) Ultrastructural identification of neuropeptides in the central nervous system. J Electron Microsc Tech 15 (1):67–80. doi:10.1002/jemt.1060150107

Margolis EB, Toy B, Himmels P, Morales M, Fields HL (2012) Identification of rat ventral tegmental area GABAergic neurons. PloS one 7 (7):e42365. doi:10.1371/journal.pone.0042365

Matsumoto M, Hikosaka O (2009) Two types of dopamine neuron distinctly convey positive and negative motivational signals. Nature 459 (7248):837–841. doi:nature08028 [pii] 10.1038/nature08028

McRitchie DA, Halliday GM, Cartwright H (1995) Quantitative analysis of the variability of substantia nigra pigmented cell clusters in the human. Neuroscience 68:539–551

McRitchie DA, Hardman CD, Halliday GM (1996) Cytoarchitectural distribution of calcium binding proteins in midbrain dopaminergic regions of rats and humans. Journal of Comparative Neurology 364 (1):121–150

Menegas W, Akiti K, Amo R, Uchida N, Watabe-Uchida M (2018) Dopamine neurons projecting to the posterior striatum reinforce avoidance of threatening stimuli. Nature neuroscience 21 (10):1421–1430. doi:10.1038/s41593-018-0222-1

Menegas W, Babayan BM, Uchida N, Watabe-Uchida M (2017) Opposite initialization to novel cues in dopamine signaling in ventral and posterior striatum in mice. eLife 6. doi:ARTN e2188610.7554/eLife.21886

Mirenowicz J, Schultz W (1994) Importance of unpredictability for reward responses in primate dopamine neurons. J Neurophysiol 72:1024–1027

Morales M, Margolis EB (2017) Ventral tegmental area: cellular heterogeneity, connectivity and behaviour. Nature reviews Neuroscience 18 (2):73-85. doi:10.1038/nrn.2016.165

Nair-Roberts RG, Chatelain-Badie SD, Benson E, White-Cooper H, Bolam JP, Ungless MA (2008) Stereological estimates of dopaminergic, GABAergic and glutamatergic neurons in the ventral tegmental area, substantia nigra and retrorubral field in the rat. Neuroscience 152 (4):1024–1031

Nusbaum MP, Blitz DM, Marder E (2017) Functional consequences of neuropeptide and small-molecule co-transmission. Nature reviews Neuroscience 18 (7):389–403. doi:10.1038/nrn.2017.56

Olszewski J, Baxter D (2014) Cytoarchitecture of the Human Brainstem. 3rd, revised and extended edn. Karger, Basel

Omelchenko N, Sesack SR (2009) Ultrastructural analysis of local collaterals of rat ventral tegmental area neurons: GABA phenotype and synapses onto dopamine and GABA cells. Synapse 63 (10):895–906. doi:10.1002/syn.20668

Orozco-Cabal L, Pollandt S, Liu J, Shinnick-Gallagher P, Gallagher JP (2006) Regulation of synaptic transmission by CRF receptors. Rev Neurosci 17 (3):279–307

Pearson J, Goldstein M, Markey K, Brandeis L (1983) Human brainstem catecholamine neuronal anatomy as indicated by immunocytochemistry with antibodies to tyrosine hydroxylase. Neuroscience 8, No. 1:3–32

Persoon CM, Moro A, Nassal JP, Farina M, Broeke JH, Arora S, Dominguez N, van Weering JR, Toonen RF, Verhage M (2018) Pool size estimations for dense-core vesicles in mammalian CNS neurons. EMBO J 37 (20). doi:10.15252/embj.201899672

Peters A, Palay SL, Webster Hd (1991) The fine structure of the nervous system: neurons and their supporting cells. 3rd edn. Oxford University Press, New York

Plant TM, Terasawa, Ei, Witchel, Selma Feldman (2015) Puberty in Non-human Primates and Man. 5 Physiological Control Systems and Governing Gonadal Function Fourth Edition:1487-1536. doi:10.1016/B978-0-12-397175-3.00032-6

Poirier LJ, Giguere M, Marchand R (1983) Comparative morphology of the substantia nigra and ventral tegmental area in the monkey, cat and rat. Brain Res Bull 11:371–397

Pomrenze MB, Giovanetti SM, Maiya R, Gordon AG, Kreeger LJ, Messing RO (2019) Dissecting the Roles of GABA and Neuropeptides from Rat Central Amygdala CRF Neurons in Anxiety and Fear Learning. Cell reports 29 (1):13–21 e14. doi:10.1016/j.celrep.2019.08.083

Rincon-Cortes M, Grace AA (2017) Sex-Dependent Effects of Stress on Immobility Behavior and VTA Dopamine Neuron Activity: Modulation by Ketamine. The international journal of neuropsychopharmacology 20 (10):823–832. doi:10.1093/ijnp/pyx048

Rodaros D, Caruana DA, Amir S, Stewart J (2007) Corticotropin-releasing factor projections from limbic forebrain and paraventricular nucleus of the hypothalamus to the region of the ventral tegmental area. Neuroscience 150 (1):8–13. doi:S0306-4522(07)01098-6 [pii] 10.1016/j.neuroscience.2007.09.043

Root DH, Wang HL, Liu B, Barker DJ, Mod L, Szocsics P, Silva AC, Magloczky Z, Morales M (2016) Glutamate neurons are intermixed with midbrain dopamine neurons in nonhuman primates and humans. Scientific reports 6. doi:Artn 30615 10.1038/Srep30615

Root DH, Zhang S, Barker DJ, Miranda-Barrientos J, Liu B, Wang HL, Morales M (2018) Selective Brain Distribution and Distinctive Synaptic Architecture of Dual Glutamatergic-GABAergic Neurons. Cell reports 23 (12):3465–3479. doi:10.1016/j.celrep.2018.05.063

Steinberg EE, Gore F, Heifets BD, Taylor MD, Norville ZC, Beier KT, Foldy C, Lerner TN, Luo L, Deisseroth K, Malenka RC (2020) Amygdala-Midbrain Connections Modulate Appetitive and Aversive Learning. Neuron. doi:10.1016/j.neuron.2020.03.016

Swanson LW (1982) The projections of the ventral tegmental area and adjacent regions: A combined fluorescent retrograde tracer and immunofluorescence study in the rat. Brain Res Bull 9:321–353

Tagliaferro P, Morales M (2008) Synapses between corticotropin-releasing factor-containing axon terminals and dopaminergic neurons in the ventral tegmental area are predominantly glutamatergic. The Journal of comparative neurology 506 (4):616–626. doi:10.1002/cne.21576

Tobler PN, Dickinson A, Schultz W (2003) Coding of predicted reward omission by dopamine neurons in a conditioned inhibition paradigm. Journal of Neuroscience 23 (32):10402–10410

Tremblay ME, Riad M, Bouvier D, Murai KK, Pasquale EB, Descarries L, Doucet G (2007) Localization of EphA4 in axon terminals and dendritic spines of adult rat hippocampus. The Journal of comparative neurology 501 (5):691–702. doi:10.1002/cne.21263

Ungless MA, Singh V, Crowder TL, Yaka R, Ron D, Bonci A (2003) Corticotropin-releasing factor requires CRF binding protein to potentiate NMDA receptors via CRF receptor 2 in dopamine neurons. Neuron 39 (3):401-407

Van Bockstaele EJ, Colago EE, Valentino RJ (1996) Corticotropin-releasing factor-containing axon terminals synapse onto catecholamine dendrites and may presynaptically modulate other afferents in the rostral pole of the nucleus locus coeruleus in the rat brain. The Journal of comparative neurology 364 (3):523–534. doi:10.1002/(SICI)1096-9861(19960115)364:3<523::AID-CNE10>3.0.CO;2-Q

Van Bockstaele EJ, Colago EE, Valentino RJ (1998) Amygdaloid corticotropin-releasing factor targets locus coeruleus dendrites: substrate for the co-ordination of emotional and cognitive limbs of the stress response. Journal of neuroendocrinology 10 (10):743–757. doi:10.1046/j.1365-2826.1998.00254.x

VanBockstaele EJ, Colago EEO, Valentino RJ (1996) Corticotropin-releasing factor-containing axon terminals synapse onto catecholamine dendrites and may presynaptically modulate other afferents in the rostral pole of the nucleus locus coeruleus in the rat brain. Journal of Comparative Neurology 364 (3):523–534. doi:Doi 10.1002/(Sici)1096-9861(19960115)364:3<523::Aid-Cne10>3.0.Co;2-Q

Verharen JPH, Zhu Y, Lammel S (2020) Aversion hot spots in the dopamine system. Curr Opin Neurobiol 64:46–52. doi:10.1016/j.conb.2020.02.002

Wanat MJ, Bonci A, Phillips PE (2013) CRF acts in the midbrain to attenuate accumbens dopamine release to rewards but not their predictors. Nature neuroscience 16 (4):383–385. doi:10.1038/nn.3335

Wanat MJ, Hopf FW, Stuber GD, Phillips PE, Bonci A (2008) Corticotropin-releasing factor increases mouse ventral tegmental area dopamine neuron firing through a protein kinase C-dependent enhancement of Ih. The Journal of physiology 586 (8):2157–2170. doi:10.1113/jphysiol.2007.150078

Wang B, Shaham Y, Zitzman D, Azari S, Wise RA, You ZB (2005) Cocaine experience establishes control of midbrain glutamate and dopamine by corticotropin-releasing factor: a role in stress-induced relapse to drug seeking. The Journal of neuroscience: the official journal of the Society for Neuroscience 25 (22):5389–5396. doi:10.1523/JNEUROSCI.0955-05.2005

Watabe-Uchida M, Zhu L, Ogawa SK, Vamanrao A, Uchida N (2012) Whole-brain mapping of direct inputs to midbrain dopamine neurons. Neuron 74 (5):858–873. doi:10.1016/j.neuron.2012.03.017

Williams SM, Goldman-Rakic PS (1998) Widespread origin of the primate mesofrontal dopamine system. Cerebral Cortex 8 (4):321-345

Yamaguchi T, Wang HL, Morales M (2013) Glutamate neurons in the substantia nigra compacta and retrorubral field. The European journal of neuroscience 38 (11):3602–3610. doi:10.1111/ejn.12359

Yuan Y, Wu W, Chen M, Cai F, Fan C, Shen W, Sun W, Hu J (2019) Reward Inhibits Paraventricular CRH Neurons to Relieve Stress. Current biology: CB 29 (7):1243–1251 e1244. doi:10.1016/j.cub.2019.02.048

